# High-density SNP array and genome sequencing reveal signatures of selection in a divergent selection rat model for aerobic running capacity

**DOI:** 10.1101/032441

**Authors:** Yu-yu Ren, Lauren G. Koch, Steven L. Britton, Nathan R. Qi, Mary K. Treutelaar, Charles F. Burant, Jun Z. Li

**Affiliations:** Department of Human Genetics, University of Michigan. Ann Arbor, MI, United States; Department of Anesthesiology, University of Michigan. Ann Arbor, MI, United States; Department of Internal Medicine, University of Michigan. Ann Arbor, MI, United States

## Abstract

We have previously established two lines of rat for studying the functional basis of aerobic exercise capacity (AEC) and its impact on metabolic health. The two lines, high capacity runners (HCR) and low capacity runners (LCR), have been selectively bred for high and low intrinsic AEC, respectively. They were started from the same genetically heterogeneous population and have now diverged in both AEC and many other physiological measures, including weight, body composition, blood pressure, body mass index, lung capacity, lipid and glucose metabolism, and natural life span. In order to exploit this rat model to understand the genomic regions under differential selection within the two lines, we used SNP genotype and whole genome pooled sequencing data to identify signatures of selection using three different statistics: runs of homozygosity, fixation index, and aberrant allele frequency spectrum, and developed a composite score that combined the three signals. we found that several pathways (ATP transport and fatty acid metabolism) are enriched in regions under differential selection. The candidate genes and pathways under selection will be integrated with the previous mRNA expression data and future F2 QTL results for a multi-omics approach to understanding the biological basis of AEC and metabolic traits.

## Introduction

Aerobic exercise capacity (AEC) can influence many complex traits including obesity and Type 2 diabetes. We established two rat lines by divergent selection of intrinsic aerobic capacity. The high capacity runners (HCR) and low capacity runners (LCR) differed by ~9-fold in endurance running distance after 32 generations, and diverged in other physiological measures, including blood pressure, body mass index, lung capacity, lipid and glucose metabolism (reviewed in Koch et al. 2012). The LCR, relative to the HCR, manifest numerous clinically relevant conditions, including increased susceptibility to cardiac ventricular fibrillation (Lujan et al. 2006) and hepatic steatosis (Thyfault et al. 2009). At the behavioral level the LCR score higher for dysfunctional sleep (Muncey et al. 2010), diminished behavioral strategies for coping with stress (Burghardt et al. 2011), and impaired memory and learning (Wikgren et al. 2012). In contrast, the HCR have reduced weight gain (Wisloff et al. 2005), increased resistance to the deleterious effects of a high fat diet (Noland et al. 2007, Novak et al. 2010), increased capacity for fatty acid oxidation in skeletal muscle (Lessard et al. 2009) and liver (Thyfault et al. 2009), and an 28-45% higher lifespan (Koch et al. 2011). At generation 10 of selection, HCR and LCR were phenotyped across several physiological measures to test if disease features had segregated differentially between the lines. Wisloff et al. discovered that adult LCR rats develop cardiovascular risks consistent with the metabolic syndrome, including large gains in visceral adiposity, increased blood pressure, dyslipidemia, endothelial dysfunction occurring within carotid arteries, and insulin resistance (Wisloff et al. 2005). Using HCR and LCR rats from generation-18 Kivela et al. (2010) found that gene expression differences related to oxidative phosphorylation and fatty acid metabolism in skeletal muscle correlated significantly with disease risk phenotypes such as physical activity levels, serum high density lipoproteins, and mitochondrial structure. Despite all of these remarkable phenotypic differences, the genetic basis linked to AEC has not been established. We hypothesize that divergent trait selection in these two lines of rats may have resulted in selective sweeps (or partial sweeps) of variants that underlie the running capacity and associated metabolic and physiological traits. In this study, we used SNP genotyping and pooled whole-genome sequencing data from rats in both lines (HCR and LCR) and two non-adjacent generations (5 and 26) to identify signatures of selection. We implemented three different statistics as well as a composite score that combines the signals from the three statistics in an attempt to uncover swept genes/pathways. The test statistics included (1) runs of homozygosity (ROH), which captures long stretches of homozygous variants that could be due to the “hitchhiking” effect near a region under positive selection; (2) fixation index (F_st_), which measures increased genetic differentiation due to divergent selection (although it could also be due to random genetic drift); and (3) aberrant allele frequency spectrum (AFS), with which a region under selection may show a local AFS that departs from the genome-wide AFS. Previous studies have proposed several methods of combining multiple selective sweep signals into a composite signal to improve the detection of true signatures of selection (Grossman et al. 2010, Utsunomiya et al. 2013, Randhawa et al. 2014). The basic rationale, as stated in Grossman et al., is that *“If each signature provides distinct information about selective sweeps, combining the signals should have greater power for localizing the source of selection than any single test.”* Our method follows the same rationale.

Using the composite score we identified genes that have diverged between time points during selection or between the two lines, and interpreted their potential roles in relation to the trait under selection by analyzing pathway enrichment of the genes. The results therefore provide useful insight into the underlying genetic basis of intrinsic aerobic capacity and other metabolic and physiological phenotypes in our HCR-LCR rat model.

## Materials and Methods

### Study overview

The protocols of animal maintenance, phenotyping, and rotational breeding have been described previously (Koch and Britton 2001). The characterization of the genetic structure, heritability, linkage equilibrium using both pedigree and genotype data have also been published (Ren et al. 2013). This study was approved by the University Committee on Use and Care of Animals, Ann Arbor, Michigan (Approval Numbers: #08905 and #03797). The proposed animal use procedures are in compliance with University guidelines, and State and Federal regulations.

### Genotyping data collection and quality control

In this study we analyzed samples representing four sample groups: two lines (HCR and LCR) in two non-adjacent generations (G5 and G26, counting from the start of the selection experiments). We first collected genotype data for 10-12 breeders for each of the four groups. Genomic DNA was extracted from frozen liver tissue, and genotyped across 803,484 SNP loci using a SNP array described before (Baud et al. 2014). Attempts to extract DNA from generations earlier than G5 revealed that many samples in G0 and G4 were degraded. We therefore chose G5 as the earliest generation in our analysis due to its assured DNA quality. During data QC we removed 21,295 SNPs with genotype missing rate >10%. This step led to 782,189 “pass QC” SNPs, which were used in calculations of runs of homozygosity (ROH) and fixation index (F_st_), described below.

### Pooled whole-genome sequencing (WGS) and quality control

DNA for four sample groups, containing 10 female breeders from each the four groups (total n=40; 10 samples overlapped with those genotyped on the array [H5 n=2, H26 n=1, L5 n=4, L26 n=3]), was extracted from frozen liver tissue and sequenced for the whole-genome in four pools using the Illumina Hiseq system at the U-M DNA Sequencing Core. The reads were mapped to the rat reference genome RGSC-3.4 using the read alignment software *Burrows-Wheeler Alignment tool (BWA)* (Li and Durbin 2009). The average read depth for the four pools were 8.4X for HCR G5, 9.9X for HCR G26, 9.9X for LCR G5, and 9.2X for LCR G26. We made joint variant calls using *Genome Analysis Toolkit (GATK)* (McKenna et al. 2010) and obtained 8,909,190 single nucleotide variants (SNV) representing alternative alleles observed in at least one of the four pools. We considered a genotype in a given pool as missing if its read depth is less than half of the pool-average. This step led to high-quality genotypes at 6,806,440 sites for HCR G5, 7,533,943 sites for HCR G26, 7,430,142 for LCR G5, and 7,218,598 for LCR G26. In all, there are 5,101,259 SNV sites with high-quality genotypes in all four pools, and these sites were used in the identification of genomic regions with aberrant allele frequency spectrum.

### Identification of long runs of homozygosity (ROH)

ROHs for each group (HCR G5, HCR G26, LCR G5, LCR G26) were identified in *PLINK* (Purcell et al. 2007) using the pass-QC markers and the following parameters: --*homozyg-window-snp 50 --homozyg-window-missing 5 --homozyg-window-het 1 -- homozyg-window-threshold .001 --homozyg-snp 25 --homozyg-kb 500* The first parameter means that the search is by sliding a moving window of 50 SNPs across the genome to detect long contiguous runs of homozygous genotypes. An occasional genotyping error or missing genotype occurring in an otherwise unbroken homozygous segment could result in the under-calling of ROHs. To address this, we allowed five missing calls and one heterozygous call per window, as described by the second and third parameters. Each SNP is then assigned an ROH status (yes/no) based on the proportion of windows that are called homozygous among all the 50-SNP windows that overlap this SNP. We have set this proportion (the fourth parameter) to be 0.001. While the above parameters describe how to define sliding windows and SNPs as ROH or not, the ROH SNPs are then merged into longer ROH segments, with the next two parameters used to set the thresholds for the minimum number of SNPs (25) and minimum length (500 kb) needed to be called an ROH segment. ROH SNP-containing segments that are shorter than both of these two thresholds would be considered not an ROH. Note that given the SNP density of the array (about 290 SNPs per 1000 Kb), the 500 Kb threshold to be called an ROH is dominant over the 25 SNP threshold.

After finding ROHs for each animal, we calculated the ROH frequency for each of the four groups across the entire genome. Then we assigned an ROH score to each Refseq gene based on the average frequency of the ROHs that overlapped its position (a score of 1 indicates 100% of the individuals within that group have an ROH across that position). This is based on an un-weighted average of all ROH segments that overlap any part of the gene. We then calculated ΔROH for temporal (G5 vs G26) and between-line (HCR vs LCR) comparisons, for a total of four analyses. The ΔROH value for every gene was ranked, and the fractional rank was transformed into a z-score via the inverse normal distribution. The z scores are used in constructing the composite scores described below.

### Fixation index (Fst)

F_st_ is a measure of genetic differentiation between two groups. It is constructed as the squared allele frequency difference between the two groups divided by a scaling factor, such that its range is from 0 (no differentiation) to 1 (complete differentiation, i.e., the two groups are fixed for different alleles). The formula for calculating F_st_ for each SNP is

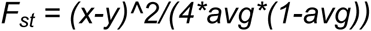

where *x* and *y* are the allele frequencies of the two groups being compared, and *avg* is the average of *x* and *y* (Wright 1965). We calculated F_st_ for every SNP for temporal (G5 vs G26) and between-line (HCR vs LCR) comparisons, for a total of four analyses (an example is shown in Figure 1). We then assigned a score to each 1 Mb window as the 80^th^ percentile of the F_st_ values of the SNPs in that window (about 290 SNPs per window). We then assigned an F_st_ value to each gene based on the unweighted average F_st_ of the windows that overlapped its position. The per-gene F_st_ values are transformed by a cubic root function when constructing the composite scores.

**Figure 1:**
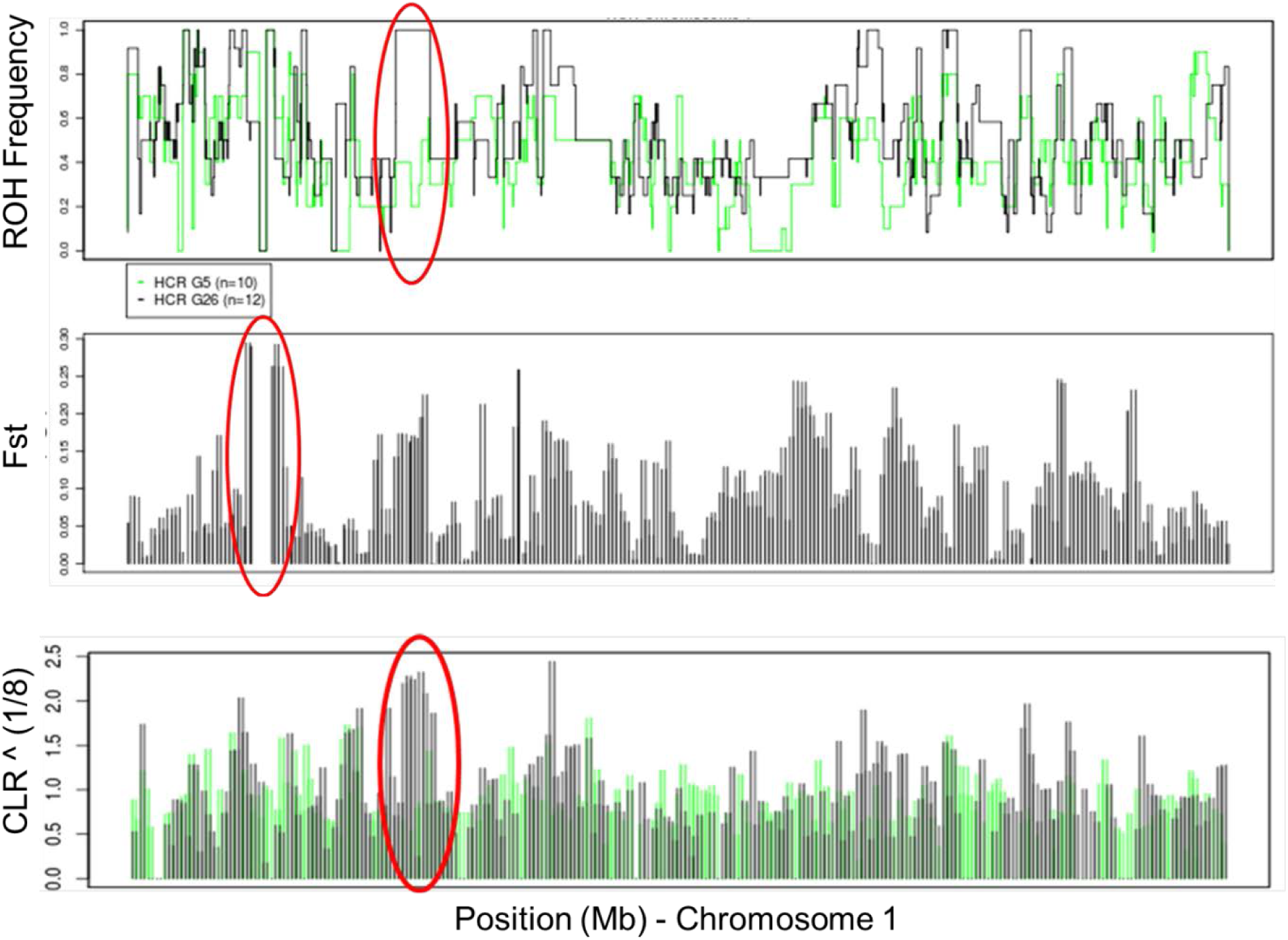
Regional examples of three selection statistics. Shown are tracks for Chromosome 1 for HCR G5 and G26. (top panel) ROH frequency intervals for HCRs G5 (green) and G26 (black). Change points are the naturally observed ROH junctions. The region highlighted in red oval represents a candidate signature of temporal selection, where all G26 animals possess a long ROH while most G5 animals do not. (middle panel) F_st_ values per 1 Mb window for the temporal comparison (G5-G26) in HCR. In the region highlighted in red oval the G5 and G26 groups have a high levels of genetic differentiation. (bottom panel) CLR values for 1 Mb windows for HCRs at G5 (green) and G26 (black). In the region highlighted in red oval the G26 animals show highly aberrant local AFS, while the G5 group does not.

### Aberrant allele frequency spectrum (AFS)

AFS analysis requires fully ascertained variant sets; that is, all variants in a region as discovered by sequencing. Genotyping data focus on pre-selected panels of SNPs and are therefore not suitable for characterizing AFS. To identify genomic regions with aberrant AFS we collected pooled WGS data, and used the observed alternative allele fraction in the pool as the surrogate of the allele frequency in each of the four groups. The difference between a local AFS, in our case defined in 1 Mb windows, and the genomewide AFS, is quantified in a parametric test described by Nielsen and colleagues, implemented in the program *SweepFinder* (Nielsen et al. 2005). When a new beneficial mutation increases in frequency in a population because of positive selection, the standing genetic variation in its neighboring region on the same chromosome will also increase in frequency (i.e., selective sweep). The pattern of allele frequencies will be skewed. *Sweepfinder* tests whether a local AFS differs from the spectrum of the whole genome by calculating a maximum composite likelihood ratio (CLR) for each window. The CLR is the ratio of the likelihood of a selective sweep to the likelihood of no sweep given the observed AFS in a window and the genome-wide AFS. It outputs the CLR statistic as well as the parameter alpha (the strength of the sweep). After calculating the CLR for each of the four sequenced pools (an example is shown in Figure 1) in 1 Mb windows, we assigned the score from the windows to genes using a non-weighted averaged of the CLR values overlapping each gene. We then calculated ΔCLR for temporal (G5 vs G26) and between-line (HCR vs LCR) comparisons, for a total of four analyses, and then transformed the ΔCLR as described below when constructing the composite scores.

### Direction of comparisons

HCR-LCR comparisons will be referred to as ‘between-line’, while the G26-G5 comparisons will be referred to as ‘temporal’. The comparisons between the two lines are bi-directional. That is, selections in either line, using the other as the reference, are meaningful and will be interpreted. For example, strong positive scores in the HCR-LCR analysis represent positive selection that occurred in the HCR line, more strongly than in the LCR. These would indicate the genomic regions under selective pressure as a consequence of the artificial selection for higher AEC. Inversely, strong negative selection score in the HCR-LCR analysis would indicate positive selection that occurred uniquely in the LCR line.

In contrast, the temporal comparisons are always G26-over-G5, as we are interested in finding signatures of positive selection at G26 using G5 as the reference. We currently do not know how to interpret these genomic regions, but it is possible that the LCR line experienced positive selection events not experienced by the HCR.

### Composite score

The main challenge of detecting signatures of positive selection is that random genetic drift could also lead to apparent peaks in certain genomic regions in ΔROH, F_st_, and ΔCLR values. As these statistics capture different aspects of the true signal, a composite score that combines the three statistics is more likely to highlight true signatures of selection above the background effects of genetic drift. In other words, regions concordant across the three test statistics will show a high composite score, whereas those with conflicting signals may show reduced composite score that is closer to the genome-wide average. Several strategies for constructing such a composite score have been described in similar studies (Grossman et al. 2010, Utsunomiya et al. 2013). In our case, each of the constituent statistics had its distinct, non-normal distribution, thus we need to transform them individually to ensure that (1) the three statistics have comparable contributions to the composite score - if the variance of one statistics is far larger than those of the other two, it will dominate the final composite score, and (2) the specificity of a scan statistics, as reflected by how frequent and how strong the peaks are, should preferably be preserved as much as possible.

We developed a novel composite score that involves transforming the three statistics with different functions.

- For ΔROH, the original scores are symmetrically distributed around 0, with a prominent peak in the middle (Figure 2a), and both tails are meaningful in that they capture the increase and decrease of ROH, respectively, in the comparison. We converted the fractional rank of ΔROH for every gene into a z-score based on the formula z = Φ^−1^(r) where Φ^−1^() is the inverse normal cumulative distribution function and r is the fractional rank, defined as n/(N+1), where n is the rank of the gene and N is the total number of genes. In the events of equal ranking; I used the function *rank(x,tie.method=“random”)* in R (R Development Core Team 2010) to add a random small noise. The resulting z score is a standard normal distribution, N(0,1), as shown in Figure 2b. The directionality is preserved after the transformation, as shown by the scatterplot (Figure 2c).
- For ΔCLR, the distribution is symmetrical around zero, with extremely strong outlier values (Figure 2d). To perform rank-based inverse normal transformation as above would have dampened the contribution of these strong peaks. To allow the peaks to make suitably large contributions to the composite score we decided to apply a cubic-root (x^1/3^) transformation. The resulting score, shown in Figure 2e, preserved the specificity of the strongest ΔCLRs. The directionality is preserved, and the majority of the observations fall near zero after the transformation (Figure 2f)
- For F_st_, the original scores are all positive, where a larger score indicates a greater population differentiation, but it does not tell which population has experienced more changes. Further, the F_st_ ‘s fall in the range of (0, 0.33) (Figure 2g), and need to be scaled up to make comparable contributions as the other two scores. To convert the one-tailed distribution into two-tailed, symmetrical distribution we attributed a sign to each F_st_ by borrowing information from the other statistics. Specifically, the assigned sign is equal to the sign of the summation of the other two transformed statistics: z-score and ΔCLR^1/3^. We then scaled up the score by a factor of 10 in order to bring the three statistics to comparable scales of variability (Figure 2h). Given that about half of the F_st_ values flipped signs, the scatterplot between the raw and transformed values is mirrored on the y-axis (Figure 2i).

**Figure 2:**
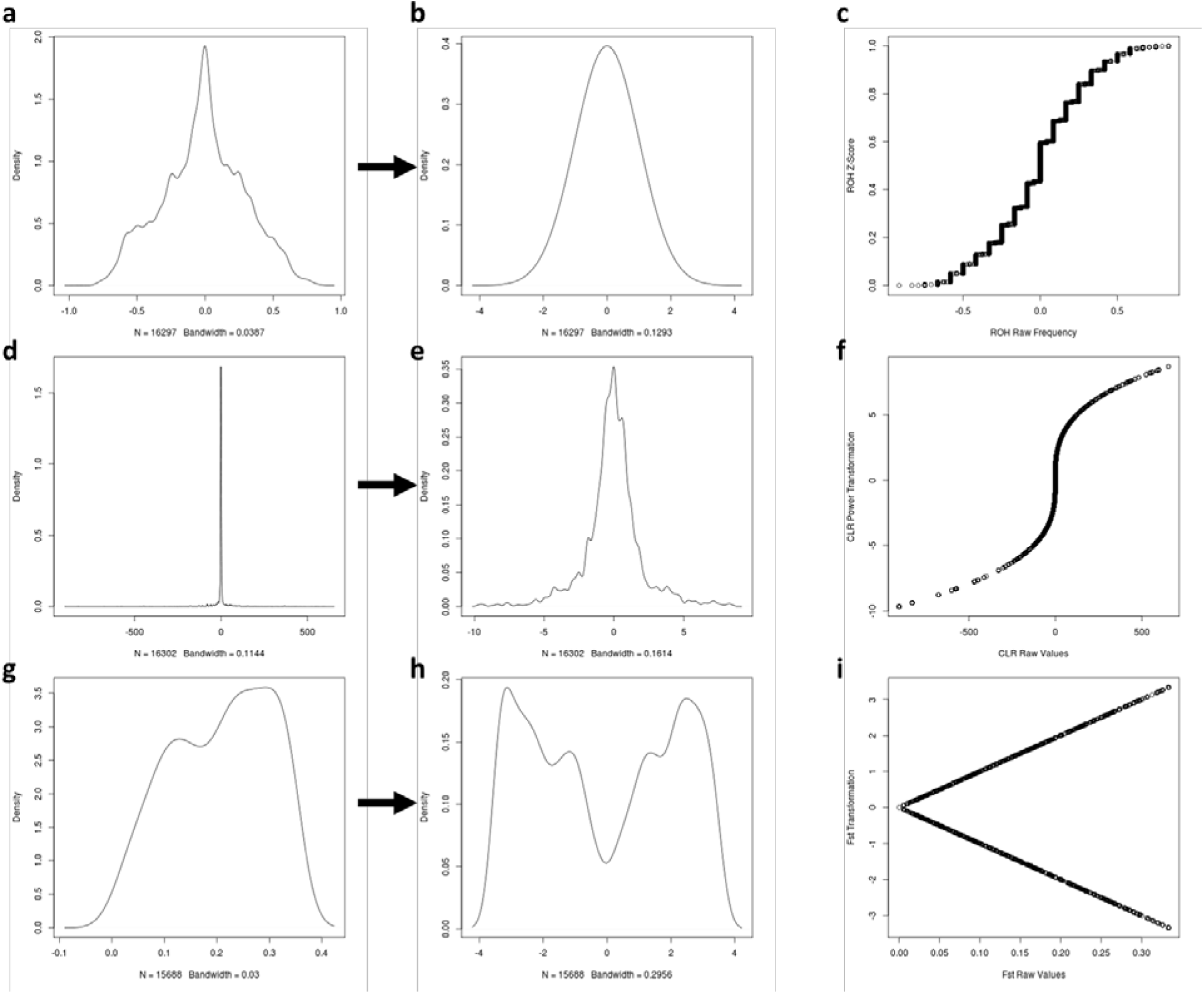
Distribution of the raw and transformed statistics. Shown are density plots for the raw values (a,d,g) and transformed values (b,e,h), and scatterplots between the raw and transformed values (c,f,i).

Finally, the composite score is the simple average of the three transformed scores. When a pseudo-p-value is needed in pathway analysis using *LRPath* (described below) we converted the composite score to the fractional rank.

### Pathway analysis

Pathway enrichment analysis was performed using *LRpath* for all genes, using the composite scores and their associated pseudo p-values. We tested for enrichment of Gene Ontology (GO) terms (across 5,948 terms) and Kyoto Encyclopedia of Genes and Genomes (KEGG) pathways (across 214 terms). The analysis was done on September 30, 2015. (Note that around June 2015 the GO annotation was updated and the results became different from those obtained before June 2015. Future updates are expected to cause further changes of the pathway analysis results. The stability of gene annotation is a nuisance in this case, but also an opportunity for the same data to return new pathway results in the future.) For a moderate correction for testing thousands of pathways we reported those that satisfied the per-pathway p-value < 0.001 for both temporal and between-line analyses.

### Visualization of enriched pathways

In many comparisons there are too many pathways that turn out to be apparently enriched and they become difficult to summarize. To facilitate the interpretation of the pathway analysis results in terms of the most salient biological signals, we needed to consolidate the top pathways into clusters of biologically related clusters, and this can be done by evaluating how any two pathways share more or fewer genes.

To detect and visualize the clustering of significant pathways from *LRpath* in an organized manner while incorporating the overlapping among gene sets, we used the *Cytoscape* plugin *EnrichmentMap (Merico et al. 2010)*. For each *LRPath* result we selected pathways passing the p-value threshold of 0.01 to build cluster maps, and defined two pathways as *connected* if they share >20% of genes between them (i.e., overlap coefficient > 0.2). In the cluster plots produced by *EnrichmentMap*, each node represents a pathway, and each green line links a pair of pathways with >20% overlap of their constituent genes. Red nodes indicate pathways showing higher composite scores in the test group than the reference group, while blue nodes indicate the opposite. The convention is that G26 and HCR are test groups and G5 and LCR are reference groups. The map of all the pairwise connections often reveal heavily connected nodes - pathways that are connected to many other pathways, thus forming the center, or “hub”, of the clusters. These clusters are highlighted by ovals; and by annotating the individual clusters we can capture the main biological signals in a given *LRPath* result. Currently there is no formal method to summarize all the pathway terms represented by a cluster as they often include very diverse concepts. We decided to apply a WordCloud algorithm (Oesper et al. 2011) that returns the most commonly used words among the pathway names in each cluster, though the resultant phrases often do not have biological meaning.

## Results

**Begin by saying more about the data if you decide to put the Methods in the end. You will need to explain what G5 is and what pass-QC SNPs are, for example. Why move Methods to the end?**

### Runs of homozygosity (ROH)

Given that selective sweeps could result in long stretches of homozygous variants, we called ROH for the two lines and two non-adjacent generations (HCR-LCR, Generations (G) 5 and 26) in *PLINK* using the ~782K pass-QC SNPs. From G5 to G26, the average number of ROH per animal did not change noticeably for HCR (383 to 382) or LCR (371 to 391); but the average length of ROH per animal increased by ~26% (3.4 to 4.3 Mb) for HCR and by ~20% (3.6 to 4.3 Mb) for LCR (Figure 3). In parallel, the average number of SNPs per ROH also increased by ~28% for HCR (981 to 1,252) and by ~19% for LCR (1,035 to 1,229) (Figure 4). Reflecting the lengthening of ROH, the average fraction of the genome covered by ROH per animal increased from 48% to 60% for HCR, and 49% to 60% for LCR; and the average gap length between ROH regions decreased by ~25% for HCR (3.2 to 2.4 Mb) and by ~27% for LCR (3.3 to 2.4 Mb) (Figure 5). We calculated ROH frequencies for each of the four groups of samples, and ΔROH for each of the four pairwise comparisons. Lastly, the ΔROH values were assigned to individual Refseq genes based on their genome coordinates.

**Figure 3:**
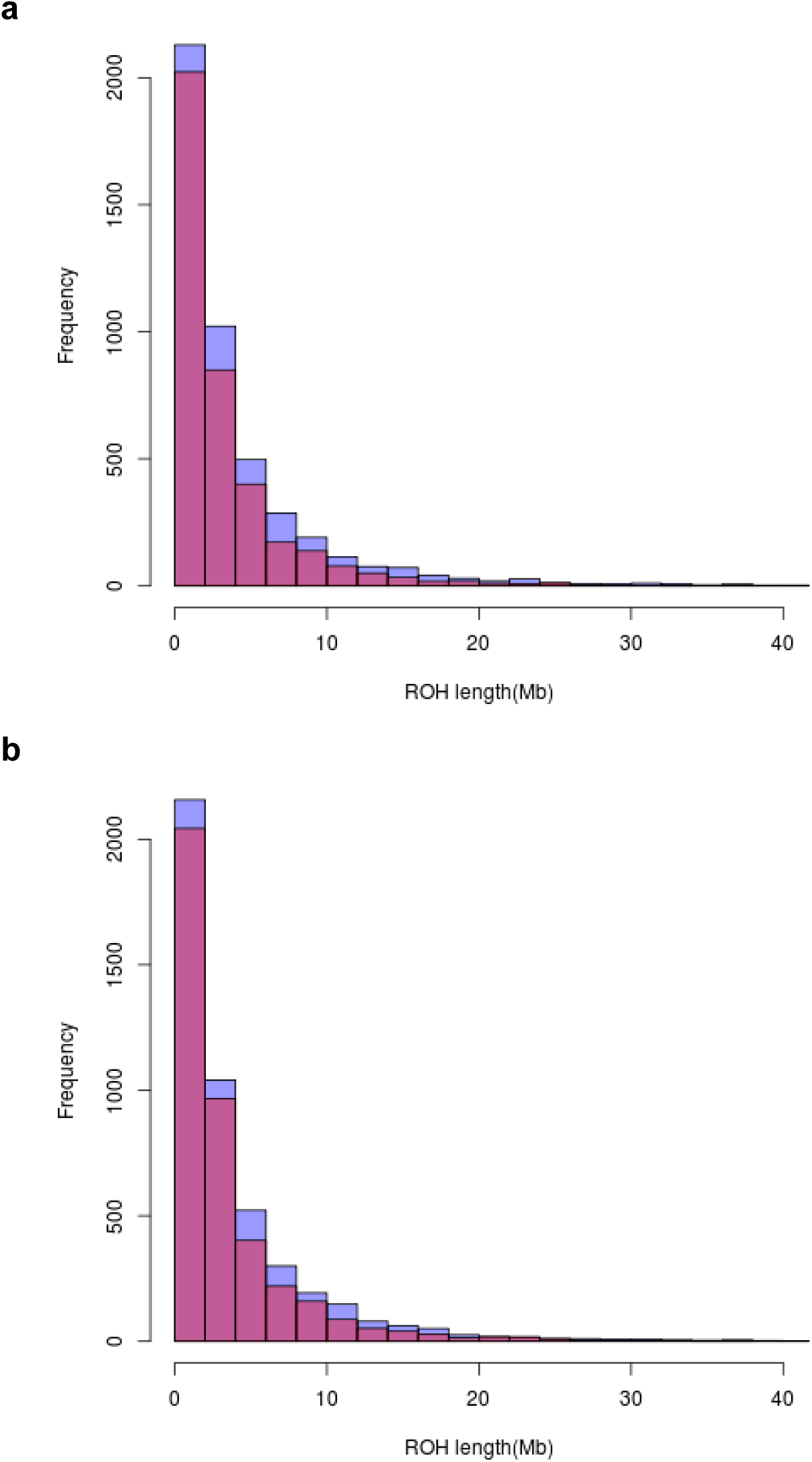
Distribution of ROH lengths in HCR (a) and LCR (b) animals.

**Figure 4:**
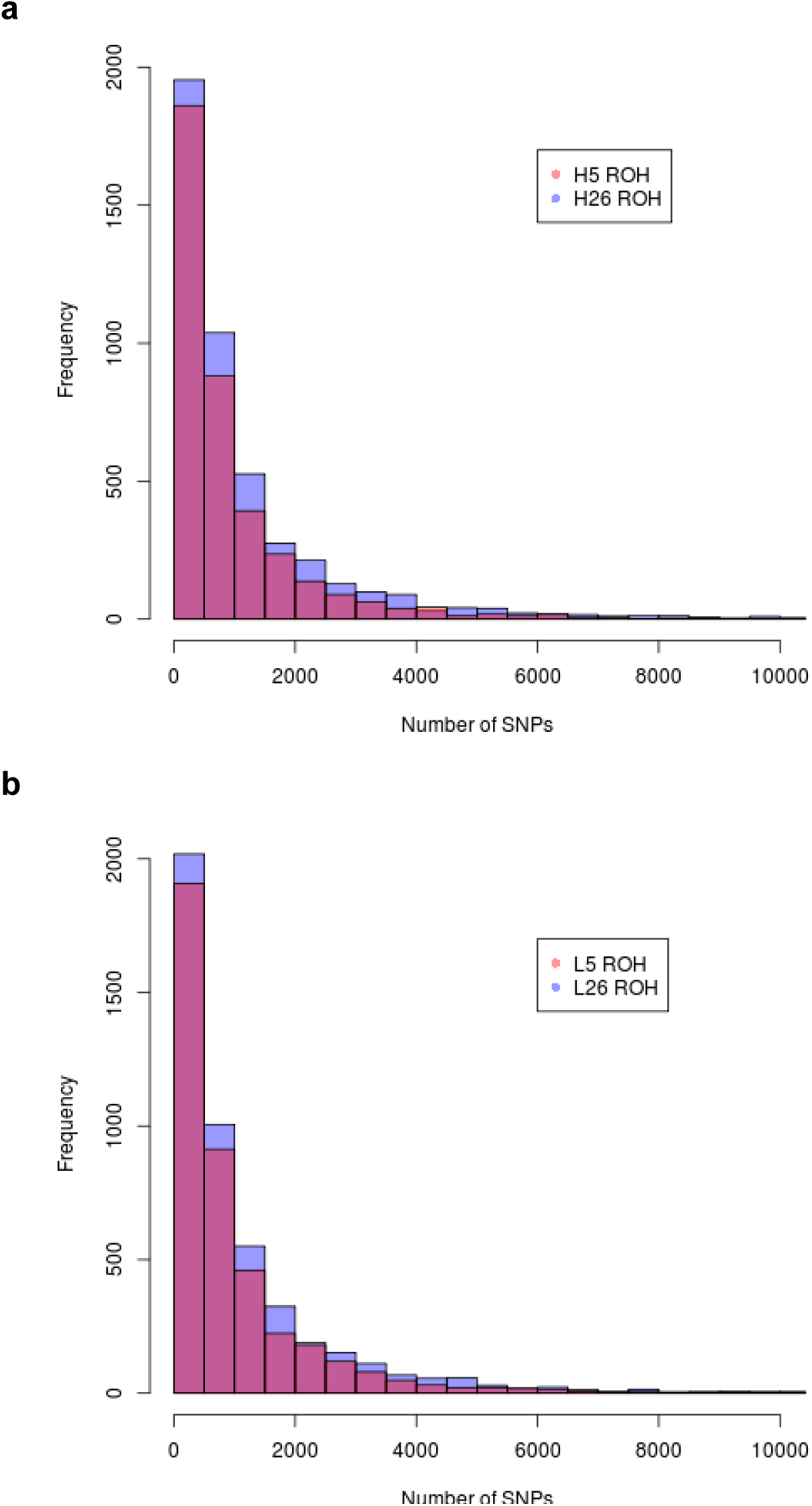
Distribution of the number of SNPs per ROH for HCR (a) and LCR (b) animals.

**Figure 5:**
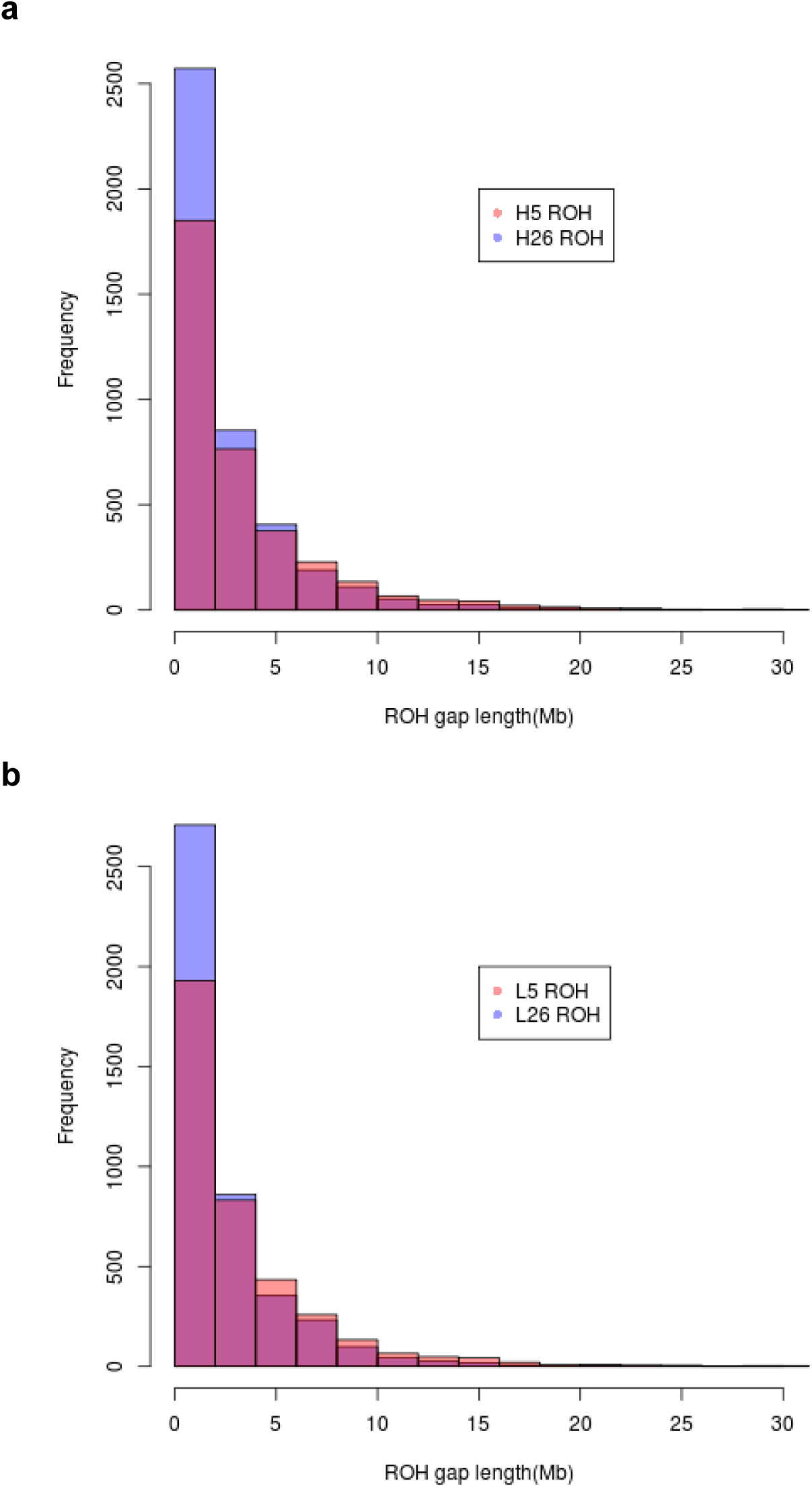
Distribution of between-ROH gap length for HCR (a) and LCR (b) animals.

### Fixation index (Fst)

Given that divergent selection often results in frequency divergence at the loci under selection, we calculated F_st_ for every SNP to determine (1) between-line allele frequency changes at both time points and (2) within-line allele frequency changes between time points, for ~782K pass-QC SNPs. The average between-line F_st_ increased from 0.038 at G5 to 0.081 at G26, while the temporal F_st_ is ~0.048 in both lines (Figure 6). After calculating F_st_ for each of the four pairwise comparisons and summariziing to 1 Mb windows we assigned F_st_ values to Refseq genes.

**Figure 6:**
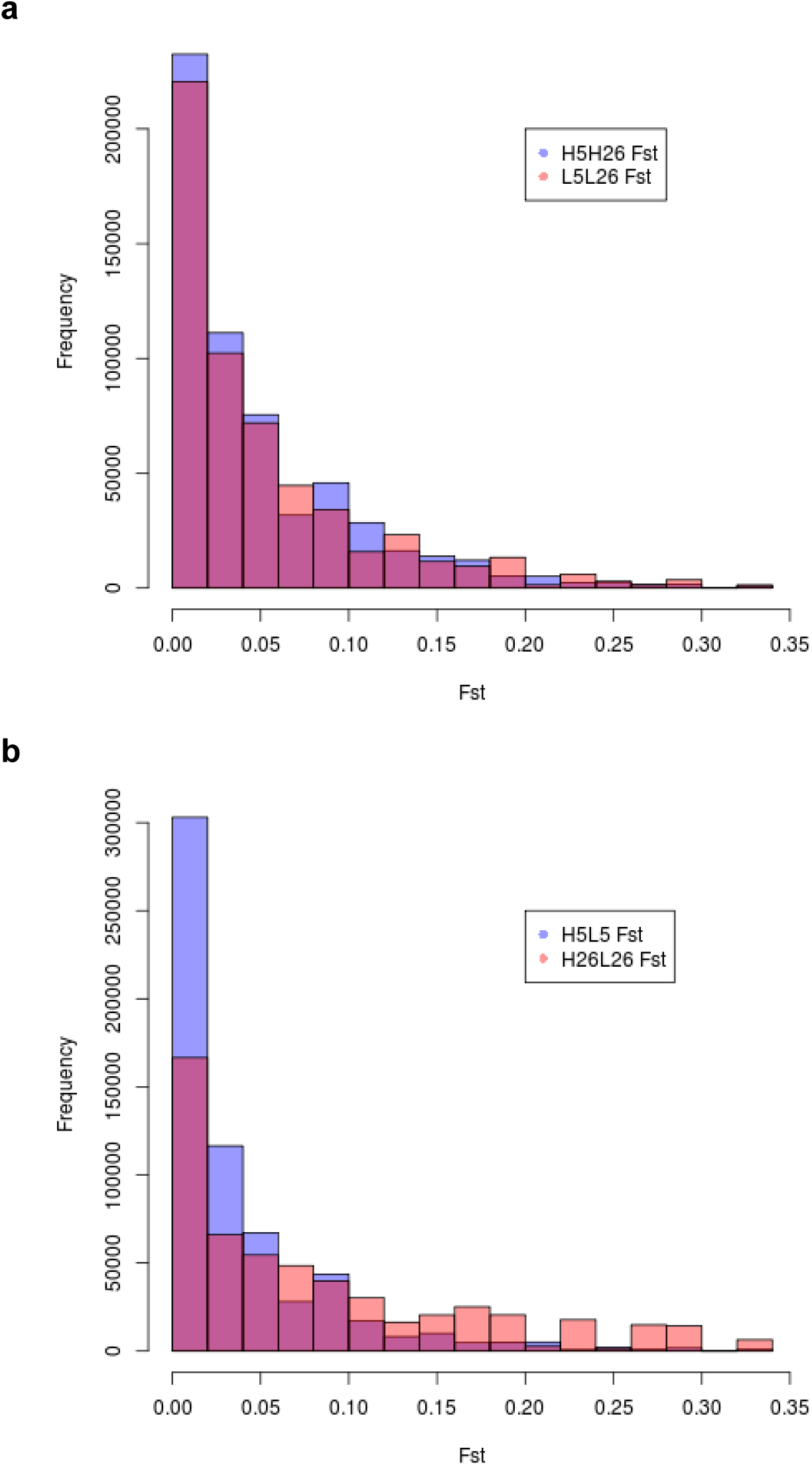
Distribution of per-SNP F_st_ values for temporal (a) and between-line (b) comparisons.

### Aberrant allele frequency spectrum (AFS)

The AFS in a region under selection may depart from that of the genomewide average. For every 1Mb window across the genome we used the high quality SNVs called from the whole-genome sequence data for the four DNA pools to calculate the composite likelihood ratio (CLR), defined as the ratio between the likelihood of selection and that of no selection given the local AFS. The average CLR increased from G5 to G26 for both lines (9.9 to 23.7 for HCR and 9.1 to 19.9 for LCR), suggesting that more regions under selection became apparent in later generations (Figure 7). After calculating ΔCLR for each of the four pairwise comparisons for 1 Mb windows we assigned ΔCLR values to individual Refseq genes.

**Figure 7:**
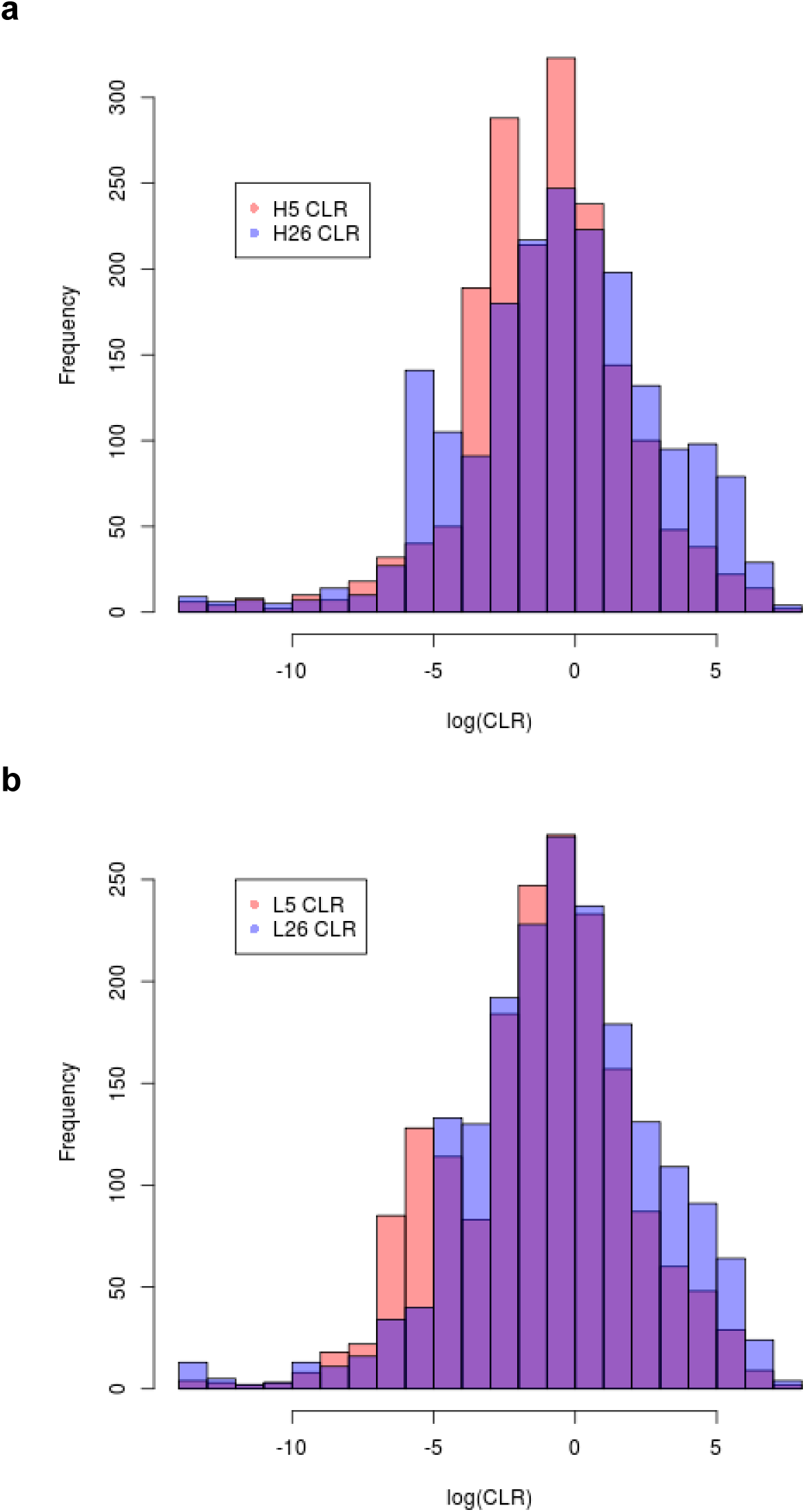
Distribution of log(CLR) values for HCR (a) and LCR (b) animals. Shown are CLR values for 1 Mb windows

### Correlations among the three scan statistics

We calculated the Spearman’s rank correlation coefficient (ρ) between each pair of pergene post-transformation test statistics (see Methods) to evaluate the level of concordance among the three signals. And we repeated the calculation for each of the four pairwise analyses, using ~16,000 genes with non-missing values in all three statistics.

For HCR G26-G5, the correlations for ΔROH - F_st_, ΔROH - ΔCLR, and F_st_ - ΔCLR are 0.61, 0.17, and 0.52, respectively (Figure 8). For LCR G26-G5, the correlations for ΔROH - F_st_, ΔROH - ΔCLR, and F_st_ - ΔCLR are 0.65, 0.22, and 0.55, respectively (Figure 9). For G5 HCR-LCR, the correlations for ΔROH - F_st_, ΔROH - ΔCLR, and F_st_ - ΔCLR are 0.59, 0.07, and 0.47, respectively (Figure 10). For G26 HCR-LCR, the correlations for ΔROH - F_st_, ΔROH - ΔCLR, and F_st_ - ΔCLR are 0.66, 0.36, and 0.63, respectively (Figure 11). Thus the recurring trend is that ΔROH and F_st_ have consistently high levels of concordance, followed by ΔCLR and F_st_. However, the concordance between ΔCLR and ΔROH is relatively low. This confirms that the three statistics are related, but they also bring distinct signatures of selection.

**Figure 8:**
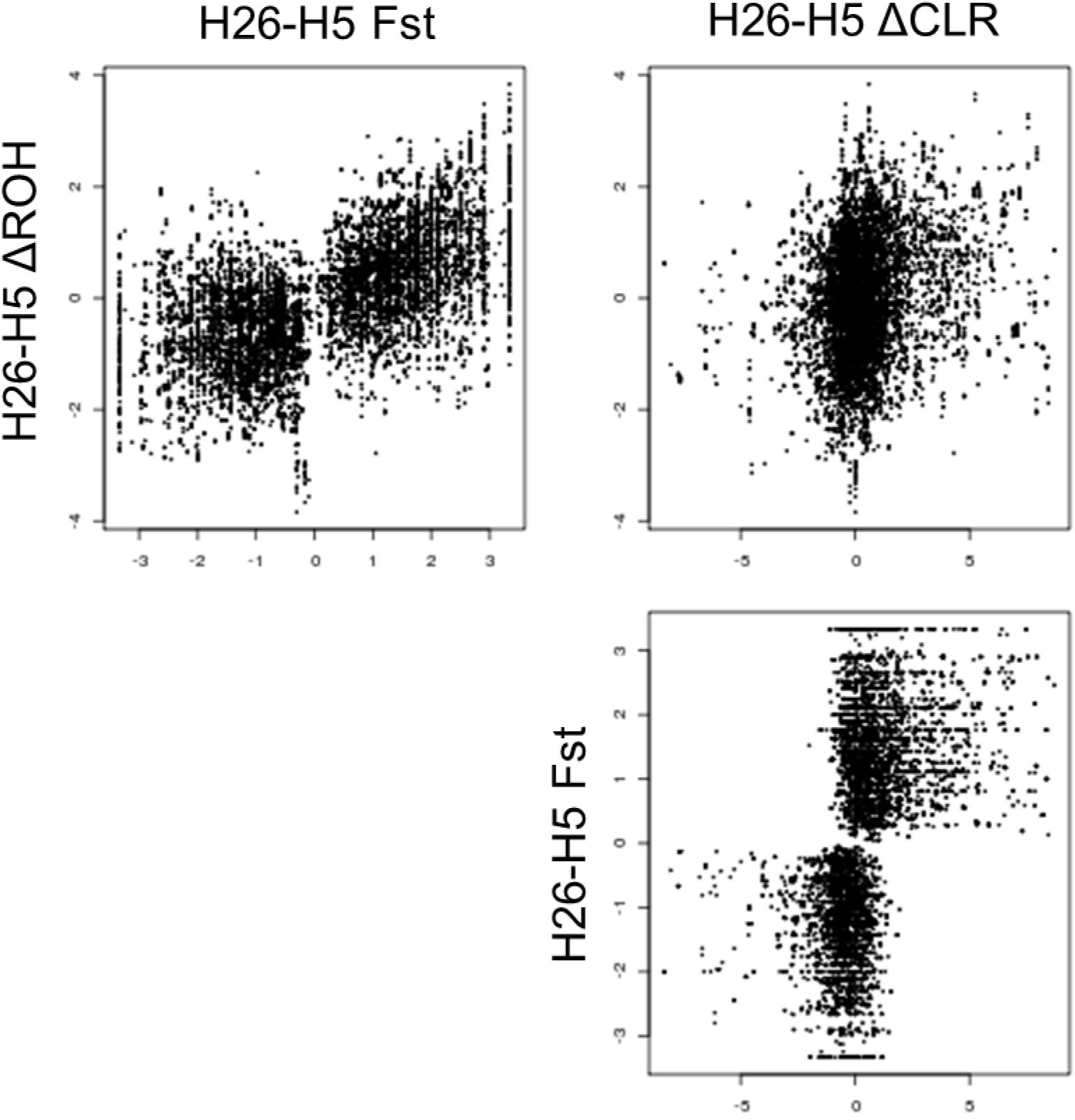
Two-way comparisons among the three test statistics for HCR G26-G5 comparisons.

**Figure 9:**
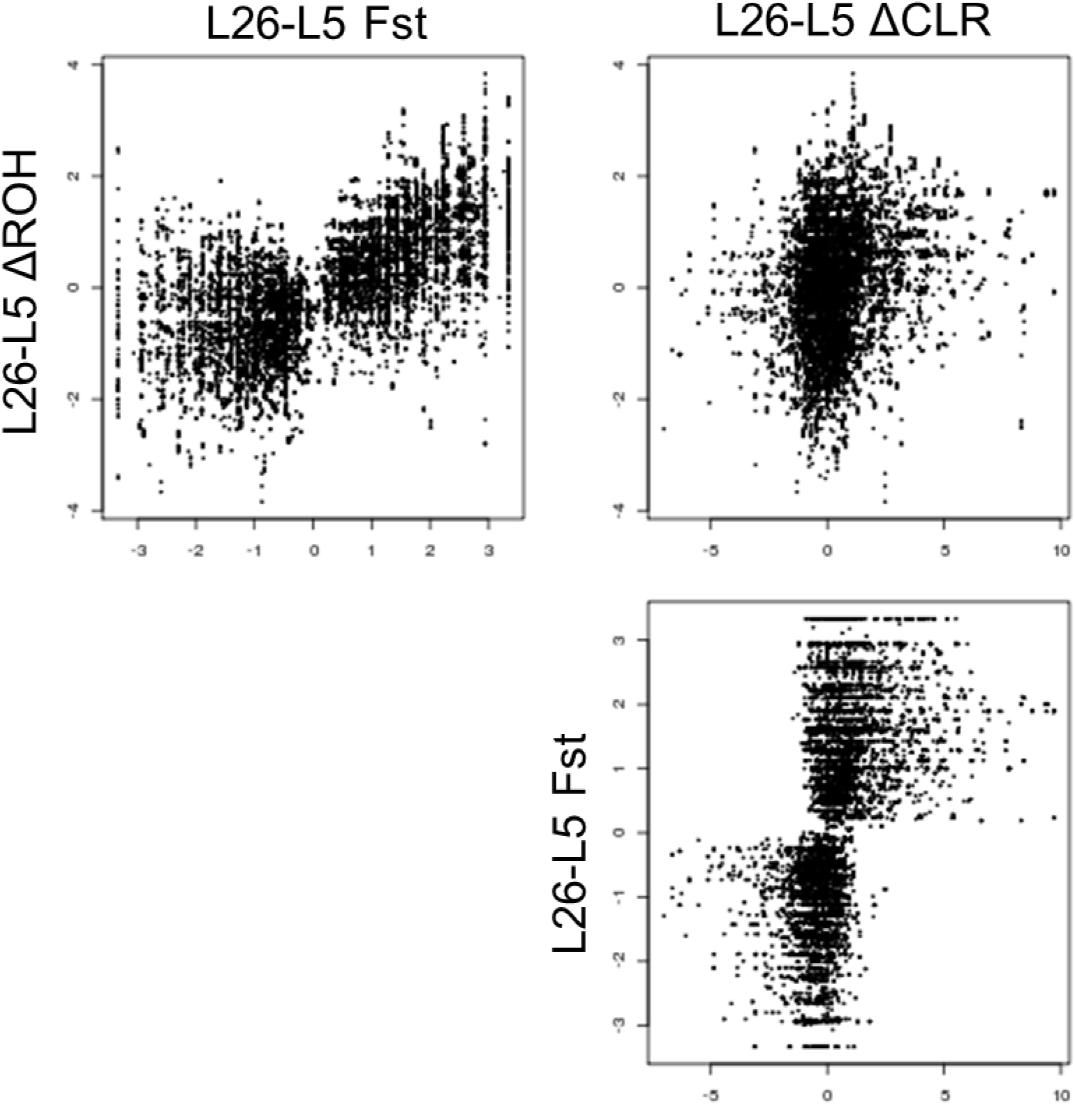
Two-way comparisons among the three test statistics for LCR G26-G5 comparisons.

**Figure 10.**
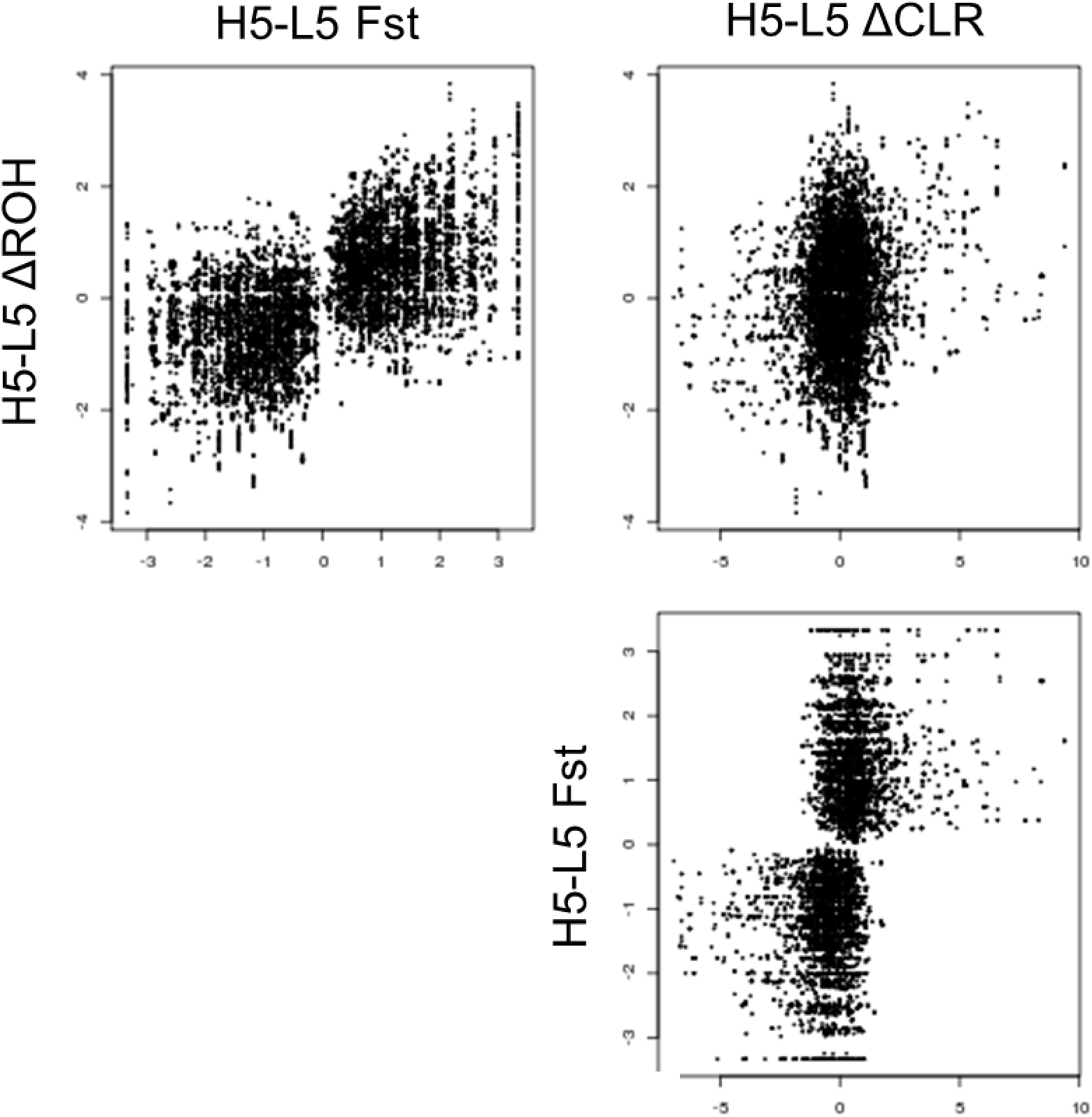
Two-way comparisons among the three test statistics for G5 HCR-LCR comparisons.

**Figure 11.**
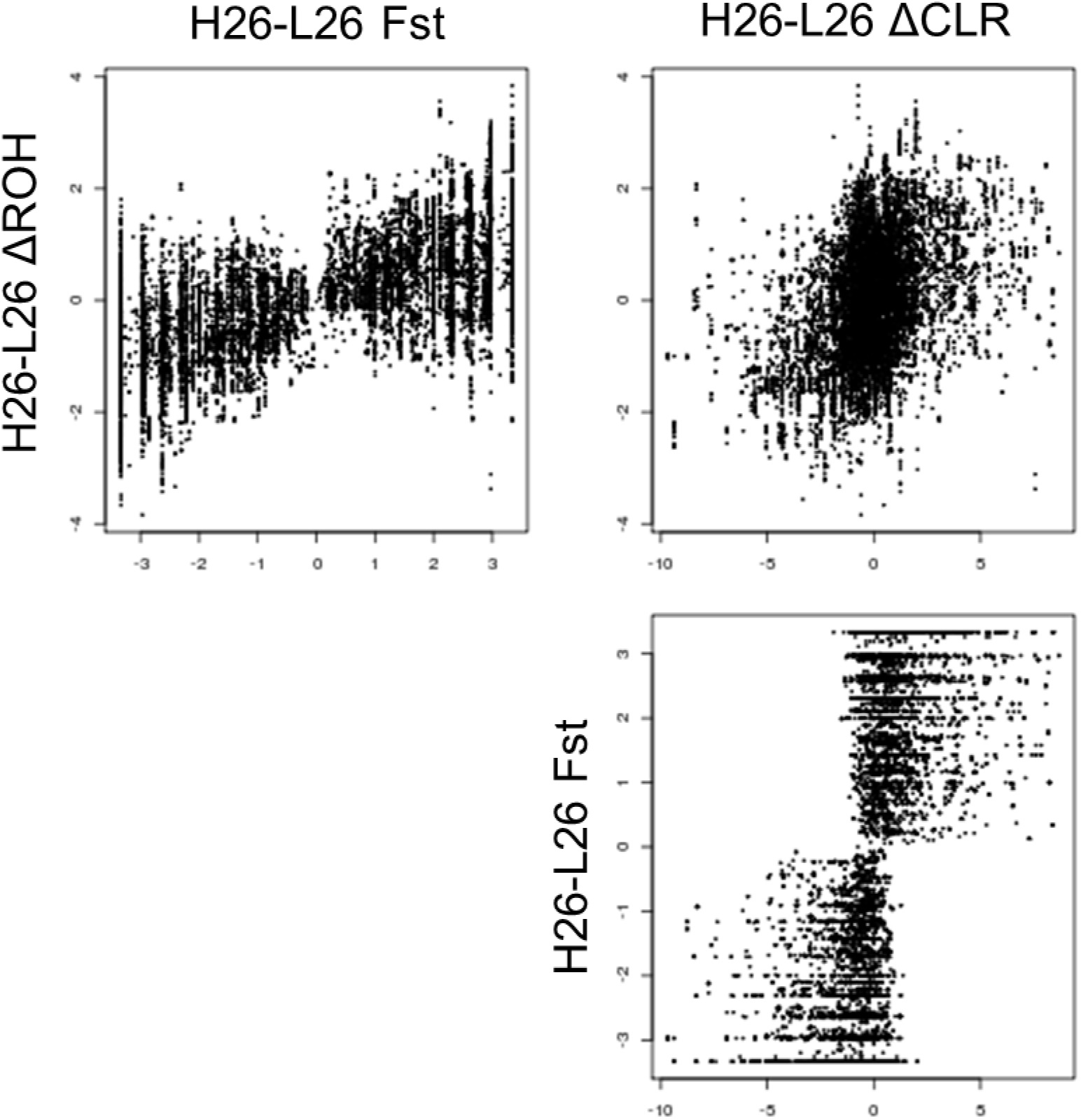
Two-way comparisons among the three test statistics for G26 HCR-LCR comparisons.

### Overlap of top ranked genes among the three statistics

To examine the genes that are highlighted in more than one statistic, we extracted the top 10% of the genes from each of the statistics and looked for overlap. This resulted in 137 genes for HCR G26-G5, 67 genes for LCR G26-G5, 79 genes for G5 HCR-LCR, and 176 genes for G26 HCR-LCR. When we evaluated pathway signals in these gene sets by using *DAVID* (Huang et al. 2007), we observed no significant pathways. In an effort to improve the power of detecting significant pathways by integrating the three statistics systematically, we developed a composite score to represent concordant selection signatures.

### Composite selection scores

We calculated the composite score to combine the three statistics described above. The composite scores for the four comparisons are shown in Figures 12. These scores do not have a natural threshold for “significance”; but given the strong phenotypic response to selection at as early as G5 (Ren et al. 2013) we expect that the genes with large HCR-LCR difference in G5 are also among the top ranked HCR-LCR genes in G26. Indeed, of the 100 genes with the highest HCR-LCR composite scores in G5, 12 also appeared among the 100 genes with the highest HCR-LCR composite scores in G26 (Table 3), while fewer than one is expected between two independent lists of 100 genes. These 12 genes fall in three top regions shared between G5 and G26 between-line composite scores: on Chromosomes 9, 16, and 18 (Figures 12c & 12d). Of particular interest are *FN1* (Fibronectin 1), which functions in cellular adhesion and extracellular matrix stability, and *PRELID2* (PRELI Domain Containing 2), which functions in phospholipid transport. These functions are relevant because we have previously found that cellular adhesion and extracellular matrix stability are two of the primary biological pathways differentially regulated in transcriptomic data as both lines age; with LCR showing greater down-regulation in both pathways as a consequence of faster aging than HCR (Ren et al. 2015). Phospholipid transport is relevant because Overmyer et al. (2015) previously found that the HCR-LCR lines differ in their fuel preference and utilization, specifically in lipids and fatty acids.

**Figure 12:**
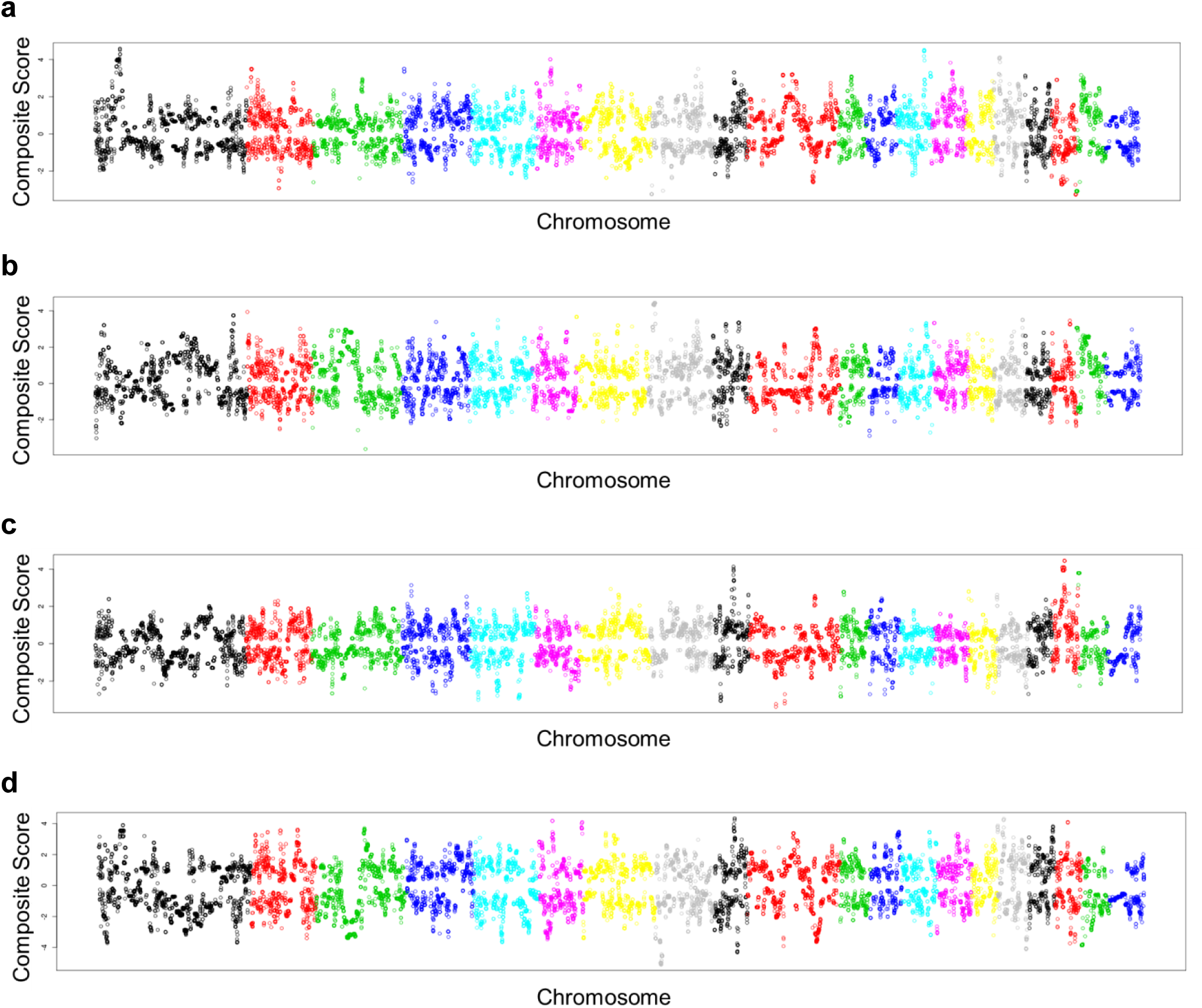
Genomewide track of per-gene composite scores. Shown are scores for HCR G26-G5 (a), LCR G26-G5 (b), G5 HCR-LCR (c), and G26 HCR-LCR (d).

We then applied the composite scores through pathway analysis. The *LRPath* algorithm returned enrichment p values for 6,162 terms and we analyzed the results in two ways.

First, we focused on the individual pathways passing p < 0.001. For temporal analyses we focused on those with higher scores in G26 than G5. In the H26-H5 analyses, the top three most significantly pathways are related to muscle contraction, including *regulation of actin filament depolymerization, myosin filament*, and *actin filament depolymerization* (Table 1). These show increased composite scors at H26, and are followed by other muscle-related pathways with slightly lower levels of significance, including *negative regulation of actin filament depolymerization, actin cytoskeleton*, and *actin filament capping*. In the L26-L5 temporal analysis there are six pathways satisfying P < 0.001 and with increased composite score at G26 (Table 2), implicating various “signaling” functions, such as *termination of signal transduction, apoptotic signaling*, and *G-protein coupled receptor signaling*. Further, for between-line analyses, the HCR-LCR comparison at G5 returned 13 pathways (Table 4), and at G26 returned 7 pathways (Table 5). As is often the case in this type of analysis, the top 3-6 pathways do not converge on 1-2 coherent functional themes. In Table 4, for example, regions with higher composite scores in HCR than LCR are enriched for genes in the *homophilic cell adhesion* pathway, echoing the aging effects found in gene expression data (Ren et al. 2015), but other pathways are difficult to interpret. Likewise, Table 5 showed that one of the top ranked pathways is for *regulation of glycogen metabolic process*, which is relevant to the metabolic differences between the two lines (Overmyer et al. 2015). There are 7 significant pathways in G5 and 4 significant pathways in G26 that are in hte HCR-low direction: these pathways show stronger signatures of selection in LCR compared to HCR (Tables 4 & 5). However, these pathways do not lead to a clear biological interpretation.

**Table 1:** Significant (p<0.0001) pathways for the composite HCR G26-G5 analysis. Only those with “up” direction are shown.

| Name | P-Value | Direction |
| --- | --- | --- |
| regulation of actin filament depolymerization | 4.44E-06 | up |
| myosin filament | 6.47E-06 | up |
| actin filament depolymerization | 7.30E-06 | up |
| regulation of protein depolymerization | 2.08E-05 | up |
| regulation of protein complex disassembly | 3.15E-05 | up |
| photoreceptor activity | 3.73E-05 | up |
| zymogen granule | 1.31E-04 | up |
| suckling behavior | 3.71E-04 | up |
| cellular response to estradiol stimulus | 4.48E-04 | up |
| zymogen granule membrane | 4.70E-04 | up |
| negative regulation of protein complex disassembly | 5.66E-04 | up |
| negative regulation of actin filament depolymerization | 6.11E-04 | up |
| actin cytoskeleton | 6.56E-04 | up |
| actin filament capping | 6.69E-04 | up |
| cytoplasmic part | 6.80E-04 | up |
| cellular response to estrogen stimulus | 7.70E-04 | up |
| protein depolymerization | 8.20E-04 | up |
| endocardial cushion morphogenesis | 8.84E-04 | up |
| protein complex disassembly | 8.93E-04 | up |
| prostate glandular acinus development | 9.15E-04 | up |

**Table 2:** Significant (p<0.0001) pathways for the composite LCR G26-G5 analysis. Only those with “up” direction are shown.

| Name | P-Value | Direction |
| --- | --- | --- |
| termination of signal transduction | 1.16E-04 | up |
| positive regulation of apoptotic signaling pathway | 1.50E-04 | up |
| termination of G-protein coupled receptor signaling pathway | 2.65E-04 | up |
| prepulse inhibition | 3.05E-04 | up |
| protein serine/threonine kinase activity | 3.43E-04 | up |
| calmodulin-dependent protein kinase activity | 5.72E-04 | up |

**Table 3:** 12 overlapping genes between the top 100 genes from the G5 and G26 HCR-LCR composite analyses.

| Gene | Name | Chromosome | Start (Mb) | End (Mb) | H5-L5 Rank | H26-L26 Rank |
| --- | --- | --- | --- | --- | --- | --- |
| SPAG16 | Sperm Associated Antigen 16 | 9 | 68.8532 | 69.7383 | 70 | 71 |
| VWC2L | Von Willebrand Factor C Domain Containing Protein 2-Like | 9 | 69.7389 | 69.9447 | 37 | 99 |
| BARD1 | BRCA1 associated RING domain 1 | 9 | 70.1202 | 70.1986 | 100 | 44 |
| ATIC | 5-Aminoimidazole-4-Carboxamide Ribonucleotide Formyltransferase/IMP Cyclohydrolase | 9 | 70.6767 | 70.6969 | 5 | 9 |
| FN1 | Fibronectin 1 | 9 | 70.7022 | 70.7712 | 81 | 93 |
| MREG | Melanoregulin | 9 | 71.3036 | 71.3580 | 37 | 28 |
| XRCC5 | X-ray repair cross-complementing protein 5 | 9 | 71.4725 | 71.5819 | 48 | 37 |
| IGFBP2 | Insulin-Like Growth Factor Binding Protein 2 | 9 | 71.9669 | 71.9942 | 97 | 47 |
| SPATA4 | Spermatogenesis Associated 4 | 16 | 0.7230 | 0.7323 | 25 | 32 |
| PRELID2 | PRELI Domain Containing 2 | 18 | 35.0600 | 35.1542 | 22 | 96 |
| GRXCR2 | Glutaredoxin, Cysteine Rich 2 | 18 | 35.1796 | 35.1941 | 28 | 68 |
| SH3RF2 | SH3 Domain Containing Ring Finger 2 | 18 | 35.2429 | 35.3445 | 55 | 19 |

**Table 4:**
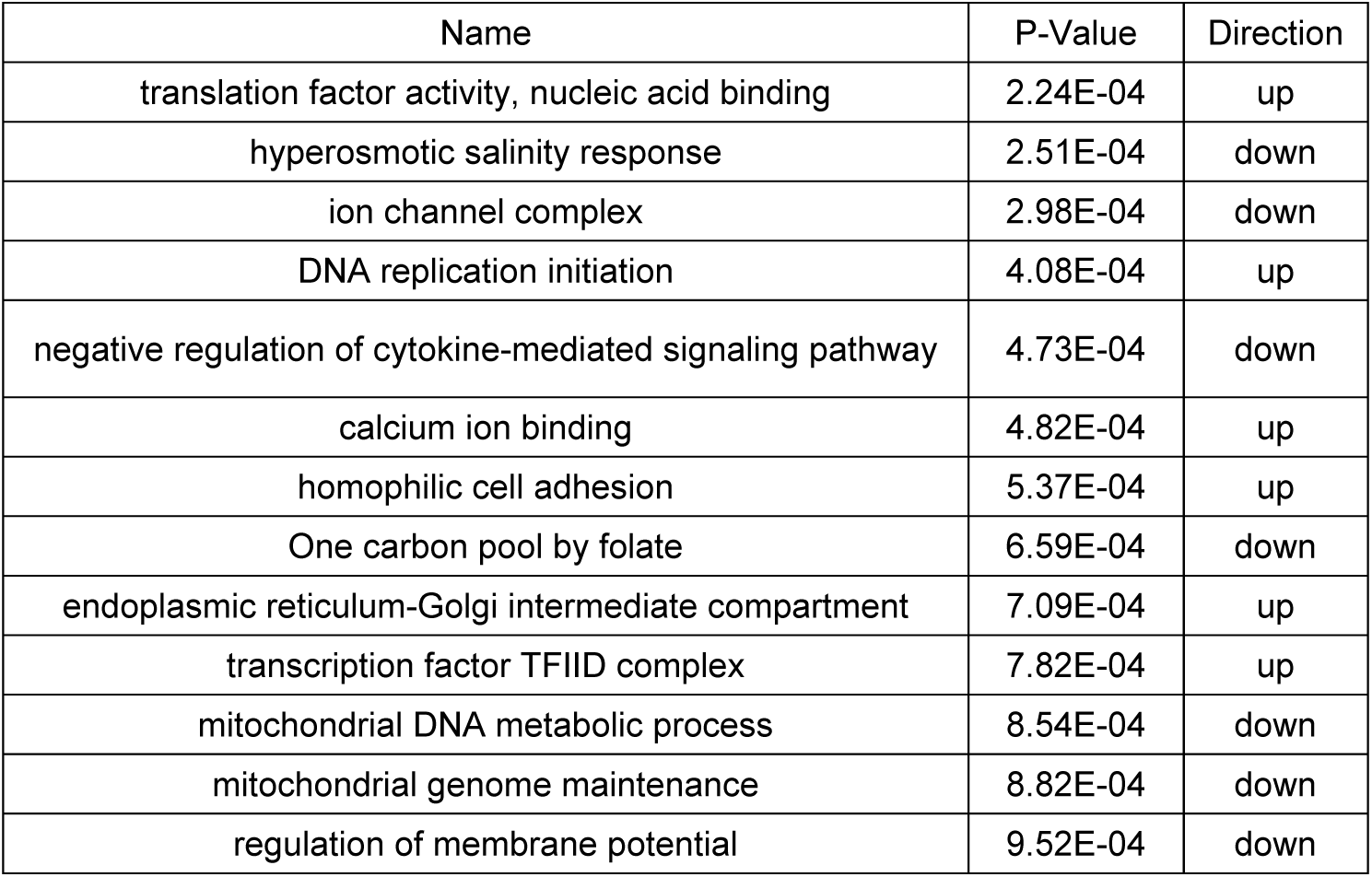
Significant (p<0.0001) pathways for composite G5 HCR-LCR analysis.

| Name | P-Value | Direction |
| --- | --- | --- |
| translation factor activity, nucleic acid binding | 2.24E-04 | up |
| hyperosmotic salinity response | 2.51E-04 | down |
| ion channel complex | 2.98E-04 | down |
| DNA replication initiation | 4.08E-04 | up |
| negative regulation of cytokine-mediated signaling pathway | 4.73E-04 | down |
| calcium ion binding | 4.82E-04 | up |
| homophilic cell adhesion | 5.37E-04 | up |
| One carbon pool by folate | 6.59E-04 | down |
| endoplasmic reticulum-Golgi intermediate compartment | 7.09E-04 | up |
| transcription factor TFIID complex | 7.82E-04 | up |
| mitochondrial DNA metabolic process | 8.54E-04 | down |
| mitochondrial genome maintenance | 8.82E-04 | down |
| regulation of membrane potential | 9.52E-04 | down |

**Table 5:** Significant (p<0.0001) pathways for composite G26 HCR-LCR analysis.

| Name | P-Value | Direction |
| --- | --- | --- |
| protein O-linked glycosylation | 1.78E-04 | up |
| cell aging | 4.17E-04 | down |
| ketone biosynthetic process | 6.20E-04 | down |
| regulation of glycogen metabolic process | 6.52E-04 | up |
| inorganic anion transport | 6.88E-04 | up |
| phototransduction, visible light | 8.98E-04 | down |
| calcium ion binding | 9.72E-04 | down |

One of the difficulties in interpreting *LRpath* results is in choosing the level of threshold: over four thousand pathways are provided in the output, each accompanied by significance levels and direction, and many of which are redundant pathways, i.e., they share the same genes with varying degrees of overlap. While in the above we examined the “significant” pathways defined at P < 0.001, our second approach is to visualize the relationship of a more relaxed set of “top” pathways, defined at P < 0.01, while taking into account the overlap of genes among the pathways. This is done by summarizing over the larger number of pathways meeting P < 0.01 using *EnrichmentMap* (see Methods).

For the temporal analysis in HCRs, there were 80 pathways and they formed many clusters (Figure 13). Two clusters on the upper right, shown in red, contain pathways that are enriched with genes with higher score in G26 than G5, with the most frequently observed words of *phospholipid-dephosphorylation* and *transport-acid-anion*, respectively (Figure 13). Note that these names do not have inherent meaning as they come from the word frequency analysis (by the *WordCloud* algorithm), thus these names are attached to the clusters to serve as provisional cluster labels. How to properly annotate the functional theme for a given pathway cluster in a formal, automated way remains a challenge. By manual annotation we determined that the *phospholipid-dephosphorylation* cluster mainly contains the pathways involved in phospholipid and fatty acid metabolism, while the *transport-acid-anion* cluster contains pathways involved in amino acid transport and metabolism. The most significant pathways related to muscle function shown in Table 1 is no longer apparent in this analysis as it involves a larger number of less significant pathways. Several other clusters in Figure 13 showed opposite direction: they contain pathways enriched with genes with higher score in G5 than G26. Such *reduced* effect of selection in later generations is difficult to interpret.

**Figure 13:**
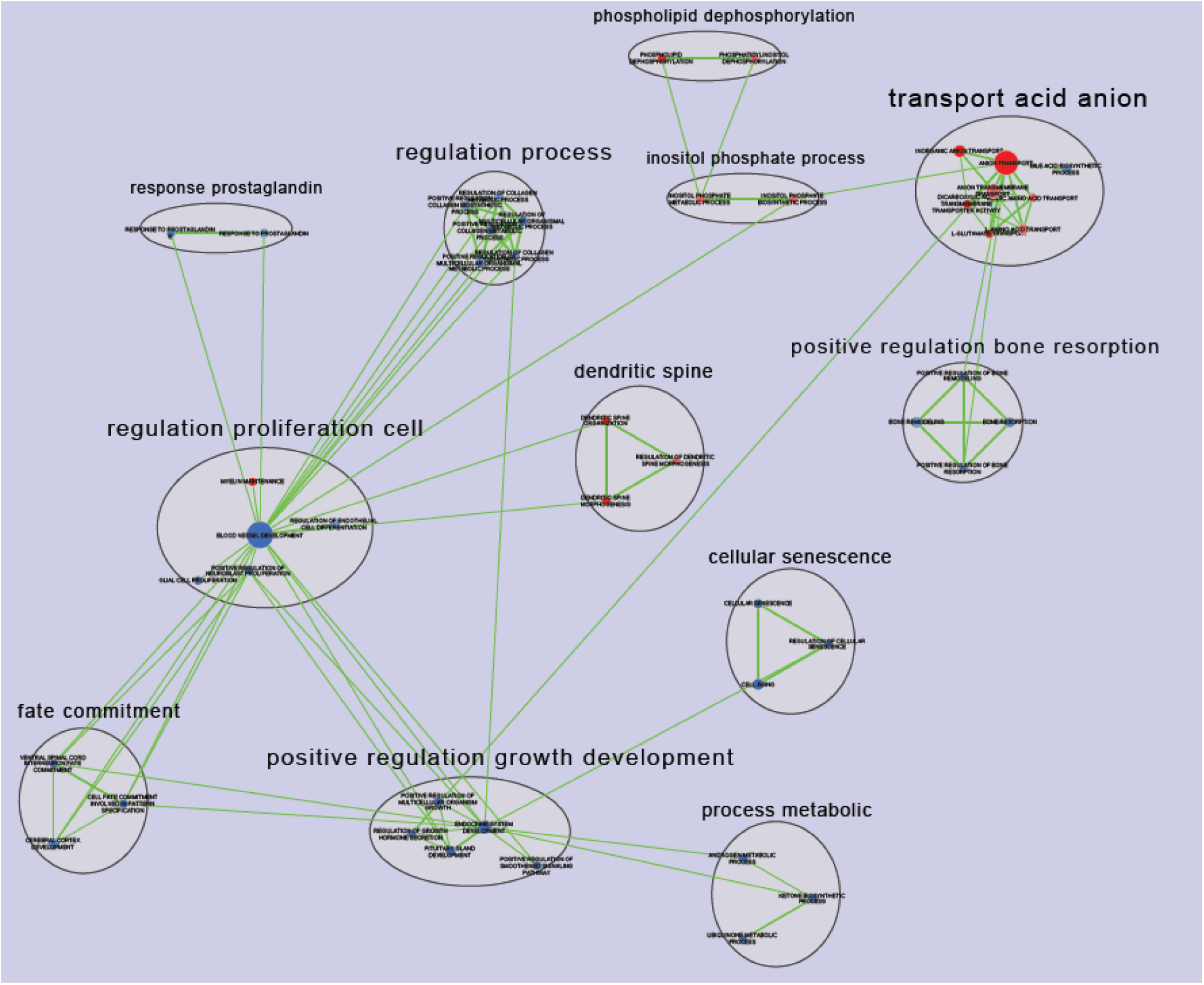
Clusters of enriched pathway for the HCR G26-G5 comparison. Shown are *EnrichmentMap* display of top pathways (P<0.01) obtained in the *LRpath* analysis of the composite scores.

For the temporal analysis of LCRs, there were 68 pathways at P < 0.01 and their clusters are shown in Figure 14. *Protein kinase* and *microtubule organization* are the major clusters enriched with genes with higher score in G26 than G5. Other clusters are difficult to interpret.

**Figure 14:**
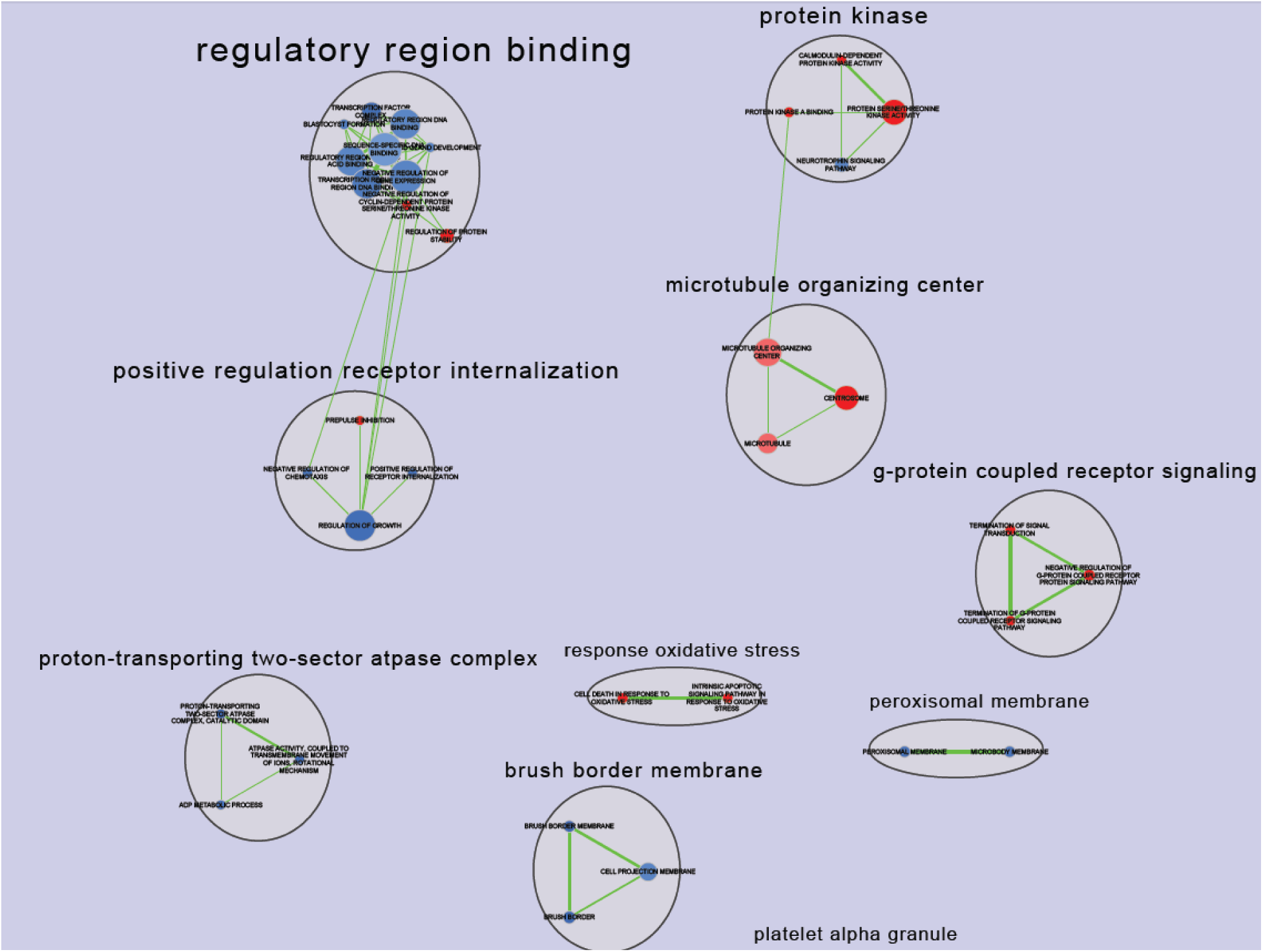
Clusters of enriched pathway for the LCR G26-G5 comparison.

The HCR-LCR comparison at G5 revealed multiple clusters containing pathways (in red) enriched with genes with higher scores in HCR than LCR, showing frequently observed words as *cytoskeletal-protein-binding, activity-transmembrane, activity-phosphatidylinositol-phospholipase*, and *growth-factor-binding* (Figure 15). By manual annotation we found that the *cytoskeletal-protein-binding* cluster contains pathways involved in actin/myosin binding and muscle contraction; the *activity-transmembrane-cluster* contains pathways involved in ATPase activity and ATP transport; the *activity-phosphatidylinositol-phospholipase* cluster contains pathways involved in phospholipid and fatty acid metabolism; and the *growth-factor-binding* cluster contains pathways involved in cellular adhesion and extracellular matrix integrity. The clusters in blue contain pathways enriched with genes with higher scores in LCR than HCR, and include frequently observed words such as *channel activity* and *phosphatase activity* (Figure 15). The *channel activity* cluster contains pathways involved in calcium and sodium channel activity; and the *phosphatase activity* cluster contains pathways involved in protein phosphatase activity.

**Figure 15:**
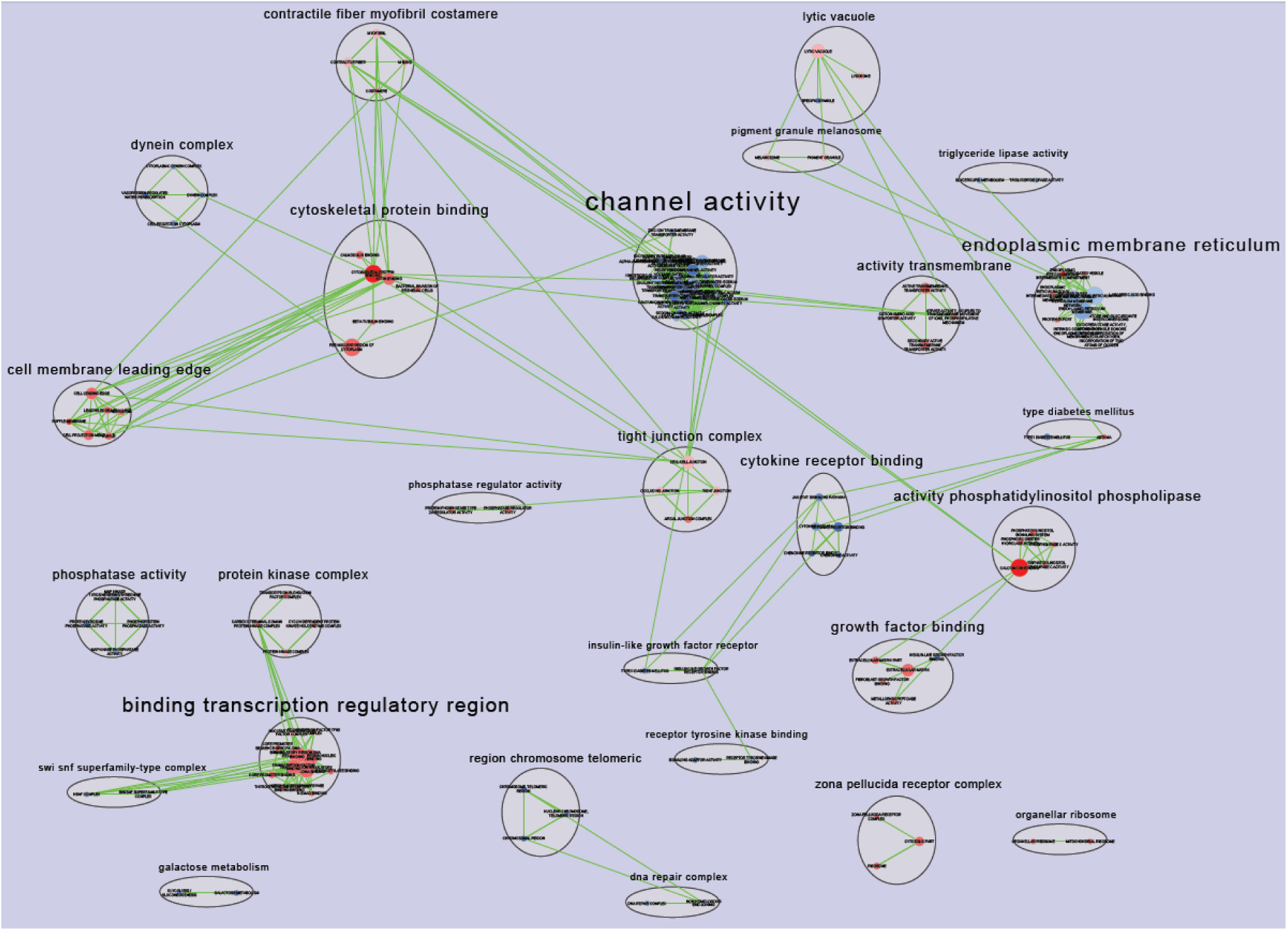
Clusters of enriched pathway for the G5 HCR-LCR comparison.

The HCR-LCR comparison at G26 (Figure 16) showed two clusters of pathways with stronger selection in HCR (in red), with words of *phospholipid-dephosphorylation* and *transport-acid-anion*, which resembles the same groups of pathways seen in the HCR G26-G5 temporal comparison (Figures 13). The clusters in blue include words such as *regulation process* and *positive regulation growth development* (Figure 16). The *regulation process* cluster contains pathways involved in the regulation of metabolic processes; and the *positive regulation growth development* cluster contains pathways involved in cellular growth and proliferation.

**Figure 16:**
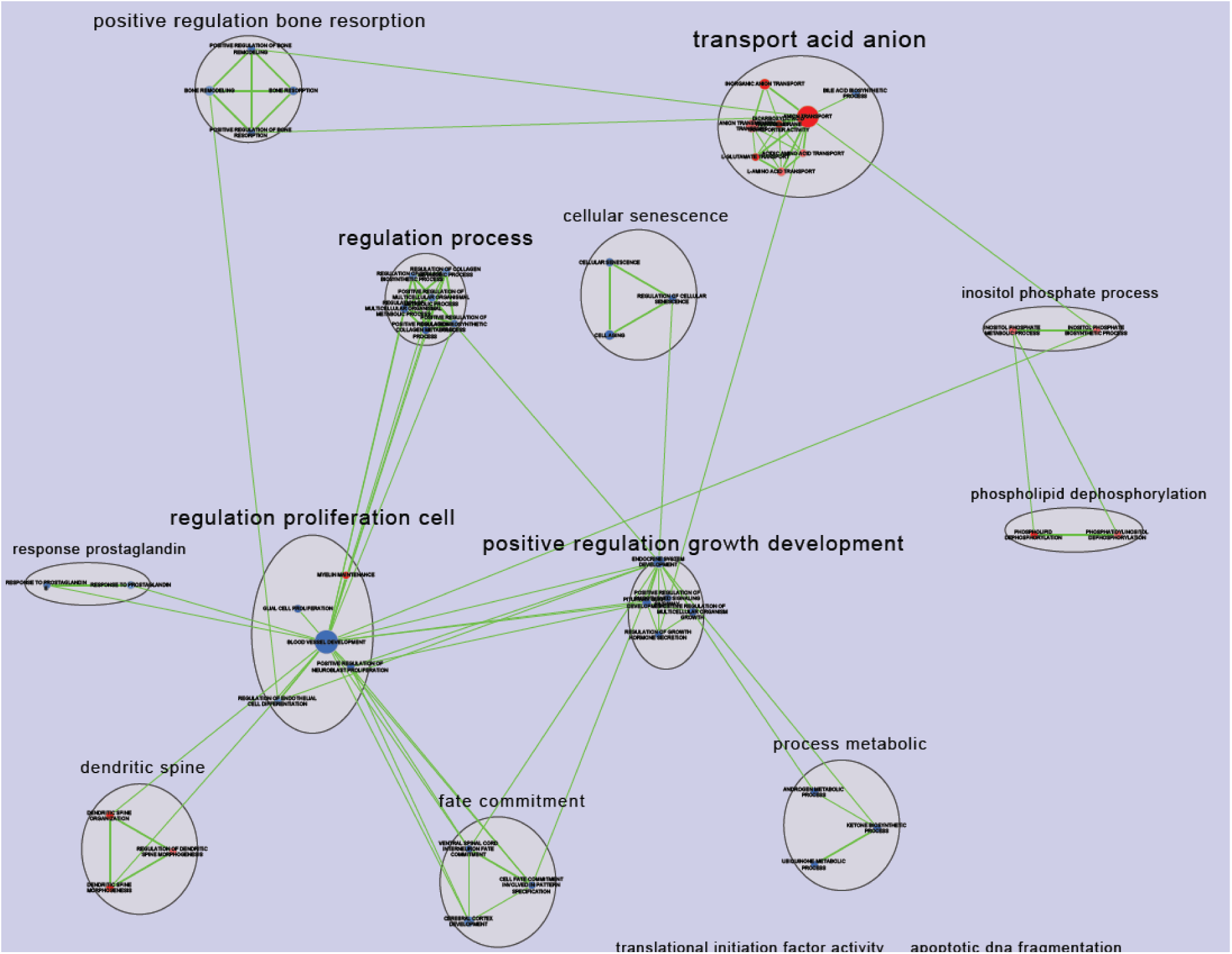
Clusters of enriched pathway for the G26 HCR-LCR comparison.

## Discussion

In this study we attempted to identify genomic regions under selection in the HCR-LCR rat model using high-density, whole-genome genotyping and pooled sequencing datasets. We identified genes and pathways under differential selection by line and by time. The majority of the increase in homozygosity between G5 and G26 in both lines is due to lengthening ROHs. The F_st_ analysis shows that the genome average of HCR-LCR differentiation increased by 2-fold from G5 and G26. The AFS analysis shows that the genome-wide CLR increased in both lines from G5 to G26, indicating the increase in aberrant local AFS and increased regions experiencing the impact of selective events at later generations.

The composite score we developed is an efficient and robust function that takes into account of the different distribution properties of the constituent test statistics. The composite signatures uncovered several physiological pathways, including those function in muscle contraction that seem to be selected in the HCR between G5 and G26, and this observation, if confirmed, offers a potential mechanistic link to the increased exercise capacity in the HCRs. In addition, the composite results provided further evidence for the importance of the aging-dependent adhesion pathways in the G5 HCR-LCR analysis.

Physical exercise is a stressful event for all complex animals. To sustain muscle contraction during exercise, the demand for adenosine triphosphate (ATP) can increase 1,000-fold compared to the resting state (Baker et al. 2010). In addition, cells must be able to have structural stability in the extracellular matrix to sustain the physiological stress. Taken as a whole, our study suggests that 1) genes in cellular integrity, actin/myosin binding and muscle contraction actin/myosin binding and muscle contraction pathways were swept in HCRs as a consequence of the increased physiological stress, and 2) genes involved in ATP-production and amino acid/lipid metabolism pathways were affected by selection and this led the HCRs to utilize multiple fuel sources to generate ATP and delay exhaustion compared to LCR during exercise. We expected that these insights into the functional consequences of selection in the HCR-LCR system provide useful information in further analysis that integrate gene expression data and direct QTL analysis in intercross populations.

## References

Baker, J. S., M. C. McCormick and R. A. Robergs (2010). “Interaction among Skeletal Muscle Metabolic Energy Systems during Intense Exercise.” Journal of Nutrition and Metabolism 2010: 13.

Baud, A., V. Guryev, O. Hummel, M. Johannesson, S. Rat Genome, C. Mapping and J. Flint (2014). “Genomes and phenomes of a population of outbred rats and its progenitors.” Sci Data 1: 140011.

Burghardt, P. R., S. B. Flagel, K. J. Burghardt, S. L. Britton, L. Gerard-Koch, S. J. Watson and H. Akil (2011). “Risk-assessment and coping strategies segregate with divergent intrinsic aerobic capacity in rats.” Neuropsychopharmacology 36(2): 390–401.

Grossman, S. R., I. Shlyakhter, E. K. Karlsson, E. H. Byrne, S. Morales, G. Frieden, E. Hostetter, E. Angelino, M. Garber, O. Zuk, E. S. Lander, S. F. Schaffner and P. C. Sabeti (2010). “A composite of multiple signals distinguishes causal variants in regions of positive selection.” Science 327(5967): 883–886.

Huang, D. W., B. T. Sherman, Q. Tan, J. Kir, D. Liu, D. Bryant, Y. Guo, R. Stephens, M. W. Baseler, H. C. Lane and R. A. Lempicki (2007). “DAVID Bioinformatics Resources: expanded annotation database and novel algorithms to better extract biology from large gene lists.” Nucleic Acids Res 35(Web Server issue): W169–175.

Kivela, R., M. Silvennoinen, M. Lehti, R. Rinnankoski-Tuikka, T. Purhonen, T. Ketola, K. Pullinen, M. Vuento, N. Mutanen, M. A. Sartor, H. Reunanen, L. G. Koch, S. L. Britton and H. Kainulainen (2010). “Gene expression centroids that link with low intrinsic aerobic exercise capacity and complex disease risk.” FASEB J 24(11): 4565–4574.

Koch, L. G., and S. L. Britton (2001). “Artificial selection for intrinsic aerobic endurance running capacity in rats.” Physiol Genomics 5(1): 45–52.

Koch, L. G., S. L. Britton and U. Wisloff (2012). “A rat model system to study complex disease risks, fitness, aging, and longevity.” Trends Cardiovasc Med 22(2): 29–34.

Koch, L. G., O. J. Kemi, N. Qi, S. X. Leng, P. Bijma, L. J. Gilligan, J. E. Wilkinson, H. Wisloff, M. A. Hoydal, N. Rolim, P. M. Abadir, E. M. van Grevenhof, G. L. Smith, C. F. Burant, O. Ellingsen, S. L. Britton and U. Wisloff (2011). “Intrinsic aerobic capacity sets a divide for aging and longevity.” Circ Res 109(10): 1162–1172.

Lessard, S. J., D. A. Rivas, Z. P. Chen, B. J. van Denderen, M. J. Watt, L. G. Koch, S. L. Britton, B. E. Kemp and J. A. Hawley (2009). “Impaired skeletal muscle beta-adrenergic activation and lipolysis are associated with whole-body insulin resistance in rats bred for low intrinsic exercise capacity.” Endocrinology 150(11): 4883–4891.

Li, H. and R. Durbin (2009). “Fast and accurate short read alignment with Burrows-Wheeler transform.” Bioinformatics 25(14): 1754–1760.

Lujan, H. L., S. L. Britton, L. G. Koch and S. E. DiCarlo (2006). “Reduced susceptibility to ventricular tachyarrhythmias in rats selectively bred for high aerobic capacity.” Am J Physiol Heart Circ Physiol 291(6): H2933–2941.

McKenna, A., M. Hanna, E. Banks, A. Sivachenko, K. Cibulskis, A. Kernytsky, K. Garimella, D. Altshuler, S. Gabriel, M. Daly and M. A. DePristo (2010). “The Genome Analysis Toolkit: a MapReduce framework for analyzing next-generation DNA sequencing data.” Genome Res 20(9): 1297–1303.

Merico, D., R. Isserlin, O. Stueker, A. Emili and G. D. Bader (2010). “Enrichment map: a network-based method for gene-set enrichment visualization and interpretation.” PLoS One 5(11): e13984.

Muncey, A. R., A. R. Saulles, L. G. Koch, S. L. Britton, H. A. Baghdoyan and R. Lydic (2010). “Disrupted sleep and delayed recovery from chronic peripheral neuropathy are distinct phenotypes in a rat model of metabolic syndrome.” Anesthesiology 113(5): 1176–1185.

Nielsen, R., S. Williamson, Y. Kim, M. J. Hubisz, A. G. Clark and C. Bustamante (2005). “Genomic scans for selective sweeps using SNP data.” Genome Res 15(11): 1566–1575.

Noland, R. C., J. P. Thyfault, S. T. Henes, B. R. Whitfield, T. L. Woodlief, J. R. Evans, J. A. Lust, S. L. Britton, L. G. Koch, R. W. Dudek, G. L. Dohm, R. N. Cortright and R. M. Lust (2007). “Artificial selection for high-capacity endurance running is protective against high-fat diet-induced insulin resistance.” Am J Physiol Endocrinol Metab 293(1): E31–41.

Novak, C. M., C. Escande, P. R. Burghardt, M. Zhang, M. T. Barbosa, E. N. Chini, S. L. Britton, L. G. Koch, H. Akil and J. A. Levine (2010). “Spontaneous activity, economy of activity, and resistance to diet-induced obesity in rats bred for high intrinsic aerobic capacity.” Horm Behav 58(3): 355–367.

Oesper, L., D. Merico, R. Isserlin and G. D. Bader (2011). “WordCloud: a Cytoscape plugin to create a visual semantic summary of networks.” Source Code Biol Med 6: 7.

Overmyer, K. A., C. R. Evans, N. R. Qi, C. E. Minogue, J. J. Carson, C. J. Chermside-Scabbo, L. G. Koch, S. L. Britton, D. J. Pagliarini, J. J. Coon and C. F. Burant (2015). “Maximal oxidative capacity during exercise is associated with skeletal muscle fuel selection and dynamic changes in mitochondrial protein acetylation.” Cell Metab 21(3): 468–478.

Purcell, S., B. Neale, K. Todd-Brown, L. Thomas, M. A. Ferreira, D. Bender, J. Maller, P. Sklar, P. I. de Bakker, M. J. Daly and P. C. Sham (2007). “PLINK: a tool set for whole-genome association and population-based linkage analyses.” Am J Hum Genet 81(3): 559–575.

R Development Core Team (2010). R: A language and environment for statistical computing. Vienna, Austria, R Foundation for Statistical Computing.

Randhawa, I. A., M. S. Khatkar, P. C. Thomson and H. W. Raadsma (2014). “Composite selection signals can localize the trait specific genomic regions in multi-breed populations of cattle and sheep.” BMC Genet 15: 34.

Ren, Y. Y., L. G. Koch, S. L. Britton, N. R. Qi, M. K. Treutelaar, C. F. Burant and J. Z. Li (2015). “Selection-, age-, and exercise-dependence of skeletal muscle gene expression patterns in a rat model of metabolic fitness.” bioRxiv doi:http://dx.doi.org/10.1101/013706.

Ren, Y. Y., K. A. Overmyer, N. R. Qi, M. K. Treutelaar, L. Heckenkamp, M. Kalahar, L. G. Koch, S. L. Britton, C. F. Burant and J. Z. Li (2013). “Genetic analysis of a rat model of aerobic capacity and metabolic fitness.” PLoS One 8(10): e77588.

Thyfault, J. P., R. S. Rector, G. M. Uptergrove, S. J. Borengasser, E. M. Morris, Y. Wei, M. J. Laye, C. F. Burant, N. R. Qi, S. E. Ridenhour, L. G. Koch, S. L. Britton and J. A. Ibdah (2009). “Rats selectively bred for low aerobic capacity have reduced hepatic mitochondrial oxidative capacity and susceptibility to hepatic steatosis and injury.” J Physiol 587(Pt 8): 1805–1816.

Utsunomiya, Y. T., A. M. Perez O’Brien, T. S. Sonstegard, C. P. Van Tassell, A. S. do Carmo, G. Meszaros, J. Solkner and J. F. Garcia (2013). “Detecting loci under recent positive selection in dairy and beef cattle by combining different genome-wide scan methods.” PLoS One 8(5): e64280.

Wikgren, J., G. G. Mertikas, P. Raussi, R. Tirkkonen, L. Ayravainen, M. Pelto-Huikko, L. G. Koch, S. L. Britton and H. Kainulainen (2012). “Selective breeding for endurance running capacity affects cognitive but not motor learning in rats.” Physiol Behav 106(2): 95–100.

Wisloff, U., S. M. Najjar, O. Ellingsen, P. M. Haram, S. Swoap, Q. Al-Share, M. Fernstrom, K. Rezaei, S. J. Lee, L. G. Koch and S. L. Britton (2005). “Cardiovascular risk factors emerge after artificial selection for low aerobic capacity.” Science 307(5708): 418–420.

Wright, S. (1965). “The Interpretation of Population-Structure by F-Statistics with Special Regard to Systems of Mating.” Evolution 19(3): 395–420.

